# Structural and functional characterisation of SiiA, an auxiliary protein from the SPI4-encoded type 1 secretion system from *Salmonella enterica*

**DOI:** 10.1101/641415

**Authors:** Peter Kirchweger, Sigrid Weiler, Claudia Egerer-Sieber, Anna-Theresa Blasl, Stefanie Hoffmann, Christiane Schmidt, Nathalie Sander, Dorothee Merker, Roman G. Gerlach, Michael Hensel, Yves A. Muller

**Author notes:** Correspondence to Yves A. Muller: Division of Biotechnology, Department of Biology, Friedrich-Alexander University Erlangen-Nuremberg, Henkestr. 91, D-91052 Erlangen, Germany.

## Abstract

*Salmonella* invasion is mediated by a concerted action of the *Salmonella* pathogenicity island 4 (SPI4)-encoded type one secretion system (T1SS) and the SPI1-encoded type three secretion system (T3SS-1). The SPI4-encoded T1SS establishes the first contact to the host membrane. It consists of five proteins (SiiABCDF) that secrete the giant adhesin SiiE. The exact mechanism by which the T1SS enables host cell recognition remains unclear. Here, we investigated structure-function relationships in SiiA, a non-canonical T1SS subunit located at the inner membrane (IM). We observe that SiiA consists of a membrane domain, an intrinsically disordered periplasmic linker region and a folded globular periplasmic domain (SiiA-PD). The crystal structure of SiiA-PD shows homology to that of MotB-PD and other peptidoglycan (PG)-binding domains. Indeed, SiiA-PD binds PG *in vitro* albeit at an acidic pH, only, whereas MotB-PD binds PG from pH 5.8 to 8. Mutation of Arg162 in SiiA impedes PG binding and reduces *Salmonella* invasion efficacy of polarized epithelial cells. SiiA forms a complex with SiiB at the IM, and the SiiA-MotB homology is likely paralleled by a SiiB-MotA homology. We show that, in addition to PG binding, the SiiAB complex translocates protons across the IM. Substituting Asp13 in SiiA impairs proton translocation. Overall, SiiA displays many properties previously observed in MotB. However, whereas the MotAB complex uses the proton motif force (PMF) to energize the bacterial flagellum, it remains to be shown how the use of the PMF by SiiAB assists T1SS function and ultimately *Salmonella* invasion.

## Introduction

*Salmonella enterica* are facultative anaerobe, Gram-negative, pathogenic bacteria that infect a wide range of hosts [1]. Infections of humans by typhoidal and nontyphoidal *S. enterica* serovars are responsible for a pleiotropy of medical conditions including intestinal inflammation and typhoid fever [2, 3]. For successful infection of polarized epithelial cells, a concerted action between the *Salmonella* pathogenicity island 4 (SPI4)-encoded type one secretion system (T1SS) and the SPI1-encoded type three secretion system (T3SS-1) is required [1, 4]. While the SPI4-T1SS establishes adhesion to the apical side of polarized epithelial cells by action of SiiE [1], the T3SS-1 mediates invasion by translocating various effector molecules (reviewed in [5]). Lack of SPI4-T1SS mediated adhesion also dramatically attenuates invasion of polarized epithelial cells by *S. enterica* serovar Typhimurium (*S*. Typhimurium) [4].

We have shown that SPI4 of *S*. Typhimurium harbours the *sii* operon encoding six proteins, namely SiiA to SiiF (SiiABCDEF) [6]. The three proteins SiiCDF form a canonical T1SS, *i.e*. SiiC corresponds to the outer membrane (OM) pore protein, SiiD to the periplasmic adapter protein and SiiF to the inner membrane (IM) ATP-binding cassette (ABC) protein. T1SS often secrete only a single protein, and the substrate of the SPI4-encoded T1SS corresponds to the largest protein present in the *Salmonella* proteome, namely the 595 kDa adhesin SiiE [6–9]. In contrast to all previous proteins, the function of the two remaining proteins SiiA and SiiB is currently only poorly understood.

It has been shown that SiiA and SiiB are integral membrane proteins that form a complex within the IM [10]. Detailed experimental data showed that the transmembrane regions of the SiiAB complex share similarities in sequence and in the topological arrangement of the membrane-spanning segments with a number of heteromeric ion-conducting channels, such as the MotAB, PomAB, ExbBD and TolQR complexes [10]. A critical aspartate residue (Asp13) in SiiA has been proposed to be located at a position homologous to Asp33 in MotB (from *S. enterica*), Asp24 in PomB (from *Vibrio alginolyticus*), Asp25 in ExbD and Asp23 in TolR (both from *E. coli*) [10–14]. ExbBD together with TonB transduce energy from the proton motive force (PMF) of the IM to high-affinity OM ion transporters [15]. MotAB, PomAB and TolQR are involved in coupling the PMF to specific protein actions at the IM. Thus, MotA and MotB (or likewise PomA and PomB) form a heteromeric complex with a 4 to 2 stoichiometry at the IM and participate in the torque generation of the flagellum of Gram-negative bacteria [12, 16]. Once activated by interaction with the C-ring of the flagellum, the complex fixes the flagellum in the membrane through the interaction of the MotB or PomB peptidoglycan (PG)-binding domain with the PG layer [16]. Torque is generated by translocating either H^+^ or Na^+^ across the IM, and the above-mentioned aspartate residues are predicted to be directly implicated in the translocation mechanism [10, 11]. The exact function of TolR is currently less well understood; overall, the Tol-Pal system plays a role in assuring the integrity of the OM [17].

The SiiA structure possibly matches the protein architecture of MotB, PomB and TolR. In the 309 residue-long MotB from *S*. Typhimurium this architecture consists of a transmembrane helix (aa 29-50), a plug helix (aa 53-66), and a C-terminal segment named ‘periplasmic region essential for mobility’ (PEM, aa 111-270) that also contains an OmpA-like PG-binding domain (aa 149-269) [18–21]. The PG-binding domain enables interaction with the PG layer in Gram-negative bacteria. This layer consists of a mesh-like structure made up of glycan strands interconnected by short peptides. The glycan strands are built from multiple molecules of N-acetylglucosamine and N-acetylmuramic acid carbohydrate molecules linked by β-1,4 glyosidic bonds. The peptide stem is a pentapeptide made up of L-Ala-D-*iso*-Glu-*meso*-A_2_pm-D-Ala-D-Ala, with *iso*-Glu being the *γ*-bonded Glu and *meso*-A_2_pm is the meso-2,6-diaminopimelic acid [22].

OmpA-like PG-binding domains occur in many proteins. They harbour a PG-binding sequence motif, *i.e*. TD-X_10_-LS-X_2_-RA-X_2-_V-X_3_-L, that is highly conserved in proteins from Gram-negative bacteria involved in PG binding, such as OmpA, MotB, PomB and PALs (PG-associated lipoproteins) [20]. Atomic insight into the structures of the OmpA-like PG-binding domain is available from many proteins; insight into the details of the interaction between PG-binding domains and PG fragments is however scarce. Positive exceptions are the crystal structure of the PG-binding domain of OmpA from *Acinetobacter baumannii* (*A. baumannii*) in complex with *meso*-A_2_pm-containing PG-derived pentapeptide as well as the NMR structure of the PG-binding domain of Pal from *Haemophilus influenza* in complex with a PG precursor [23, 24]. Although the individual functions of proteins containing PG-binding domains are as diverse as forming OM pores, involved in antibiotic and small molecules export (OmpA), or generating torque for propelling the flagellum (MotB, PomB), they all share a common domain core fold with a β1/α1/β2/α2/β3/β4 secondary structure topology [19, 25].

Here, we analysed the structure and function of SiiA from *S*. Typhimurium. The recombinantly produced periplasmic region of SiiA (ppr-SiiA, aa 40-210) was studied using structure predictions, limited proteolysis, mass spectrometry and circular dichroism (CD). The crystal structure revealed that SiiA shares structural homology with other OmpA-like PG-binding domains, among them MotB. With the help of *in vivo* and *in vitro* PG-binding studies, we observed a pH-dependent binding of SiiA to PG and unravelled a link between PG binding and *Salmonella* invasion efficacy. We also showed that the IM-located SiiAB complex pumps indeed protons across the membrane similar to what has been observed for MotAB and other members of the family of heteromeric ion-conducting channels, which use the PMF to generate work.

## Results

### Domain architecture of SiiA

Weak sequence homologies were detected between the transmembrane regions of the SiiAB proteins and MotAB, ExbBD, and TolQR, whereas no sequence homologies were initially observed for the remaining regions of SiiA and SiiB [10]. In order to gain predictive insight into the entire structure of 210 residue-long SiiA, its sequence was analysed with program XtalPred [26]. This bioinformatics analysis suggests that SiiA consists of a transmembrane (TM) helix (residues 14-38), a disordered and unfolded linker (aa 51-92) and a C-terminal globular domain that extends from residues 93 to 210.

Ppr-SiiA was purified from *E. coli* in order to experimentally validate the predicted domain architecture. When subjecting purified ppr-SiiA to a limited proteolysis experiment with the unspecific protease thermolysin then a single stable fragment of about 12 kDa is readily formed (Fig. 1a). Such a behaviour is commonly observed for proteins that consist of a folded globular domain from which disordered segments extend. Access of these segments by proteases is facilitated; hence, these segments are especially prone to proteolytic degradation (see for example [27]). We named the protease-resistant 12 kDa fragment: SiiA periplasmic domain (SiiA-PD). Its composition was analysed by mass spectrometry, and analysis of peptides obtained through tryptic digestion reveals multiple sequence coverage for SiiA residues 104 to 206 (Fig. S1a). Please note that this analysis does not allow for the unambiguous determination of the exact starting and ending residue of the protease-generated SiiA-PD fragment.

**Fig. 1.**
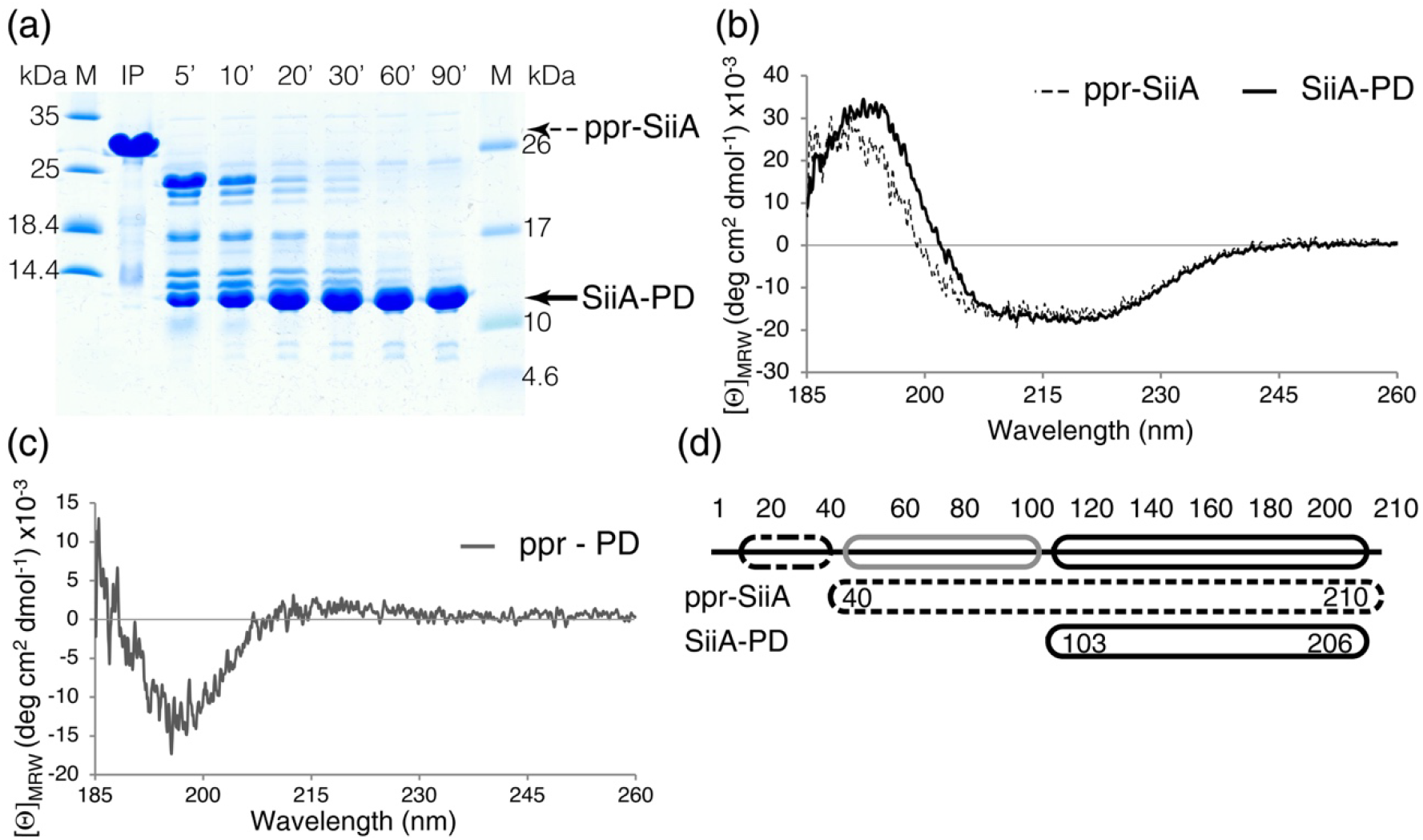
Domain structure of SiiA. (a) Limited proteolysis of ppr-SiiA (about 26 kDa) with the unspecific protease thermolysin monitored by SDS-PAGE. Limited proteolysis yields SiiA-PD (about 12 kDa), the folded C-terminal domain of SiiA. (b) CD spectrum of ppr-SiiA (after removal of the HisTag with thrombin) and SiiA-PD. (c) CD difference spectrum calculated by subtracting the spectrum of SiiA-PD from the spectrum of ppr-SiiA. (d) Schematic representation of the domain structure of SiiA, consisting of a predicted transmembrane region, an intrinsically unstructured region and a folded periplasmic domain (SiiA-PD). The existence of the latter two has been corroborated by the experimental data presented here. In (a), (b) and (d) ppr-SiiA data are displayed/marked with dashed lines and SiiA-PD with black continuous lines; in (d) the intrinsically disordered region is indicated with a gray line.

CD spectroscopy measurements were performed, and the secondary structure composition of ppr-SiiA and of SiiA-PD compared. The CD spectra show that both protein variants display spectra corresponding to folded proteins with mixed secondary structure elements (Fig. 1b). Differences in the secondary structure composition become apparent when calculating a difference CD spectrum in which the CD signal of SiiA-PD is subtracted from ppr-SiiA (Fig. 1c). The difference spectrum corresponds to the CD spectrum of an unfolded protein devoid of any secondary structure elements. This indicates that segments that are not part of SiiA-PD, but present in addition in ppr-SiiA, are likely disordered in solution. This is in line with the observation that both protein variants display nearly identical melting temperatures (T_M_) (Table S1). While SiiA-PD unfolds at 79°C, the entire periplasmic region of SiiA unfolds at 78°C showing that segments that are present in addition in ppr-SiiA do not provide any further stabilizing interactions in SiiA.

SiiA-PD and ppr-SiiA elute both as dimers in an analytical size exclusion chromatography run (Fig. S1b, Table S2). Hence, the supplemental protein segments present in ppr-SiiA do not alter the oligomerisation state of ppr-SiiA. The experiments from above show that, following the membrane domain (SiiA-MD, aa 1-39), SiiA consists of an unstructured periplasmic linker region (SiiA-LR) and a dimeric periplasmic domain (SiiA-PD) (Fig 1d).

### Crystal structure of SiiA-PD

We determined the crystal structure of SiiA-PD at 1.9 Å with the MAD method using SeMet-labelled protein (Table 1). Six molecules are present in the asymmetric unit and form three identical dimers. The terminal residues 103 and 203 are visible in all six monomers, and up to three additional C-terminal residues visible in some of the molecules. Absence of electron density for residues in the loop segment that interconnects the secondary structure elements β2 and α2 (aas 153 to 156) leads to a chain break in three monomers. When calculating an anomalous difference density map with phases derived from the refined structure then electron density is observed for all methionines present in the asymmetric unit (Table S3) thereby corroborating model building and structure refinement.

**Table 1.** Crystallographic data collection, phasing and refinement statistics.

| PDB deposition code | 6QVP |  |  |
| --- | --- | --- | --- |
| **Data collection** |  |  |  |
| Space group | P1 |  |  |
| Cell dimensions |  |  |  |
| *a*, *b*, *c* (Å) | 58.33, 58.36, 65.83 |  |  |
| $\alpha$, $\beta$, $\gamma$ (°) | 93.9, 94.0, 119.00 | | |
|  | Peak | Inflection | Native |
| Wavelength (Å) | 0.9797 | 0.9799 | 0.9184 |
| Resolution (Å) | 42.0 – 1.9<br>(2.01 - 1.9)<sup>b</sup> | 42.5 - 1.94<br>(2.05 – 1.94) | 42.5 – 1.9<br>(2.01-1.90) |
| *R*<sub>meas</sub> (%) | 12.0 (62.6) | 9.2 (58.9) | 11.4 (54.4)<sup>c</sup> |
| *I*/$\sigma$(*I*) | 7.1 (2.0) | 8.6 (1.9) | 7.5 (2.3) |
| CC<sub>1/2</sub> (%) | 99.1 (67.1) | 99.5 (75.1) | 99.1 (72.6)<sup>c</sup> |
| Resolution limit anomalous signal (Å)<sup>d</sup> | 2.4 | 3.9 | 4.7 |
| Completeness (%) | 95.9 (93.7) | 95.5 (93.6) | 95.9 (93.8) |
| Redundancy | 1.7 | 1.7 | 1.7 |
| **Refinement** |  |  |  |
| Resolution range (Å) |  |  | 19.9 – 1.9 |
| No. of unique reflections |  |  | 57795 |
| *R*<sub>work</sub>/*R*<sub>free</sub> |  |  | 18.6/24.6 |
| Ramachandran (%) |  |  |  |
| favored/outlier |  |  | 97.6/0.0 |
| Total no. of atoms |  |  | 5518 |
| No. of protein atoms |  |  | 4899 |
| No. of water atoms No. of ligand atoms |  |  | 619<br>15 |
| *B*-factors (Å)<sup>2</sup> |  |  |  |
| Protein |  |  | 35.3 |
| Water |  |  | 33.9 |
| Ligands |  |  | 42.0 |
| R.m.s deviations |  |  |  |
| Bond lengths (Å) |  |  | 0.013 |
| Bond angles (°) |  |  | 1.182 |

<sup>a</sup> A single crystal was used for phasing and refinement.
<sup>b</sup> Values in parentheses refer to the highest resolution shell.
<sup>c</sup> Values calculated with the Friedel's law equal false setting. Prior to refinement, however, the Friedel pairs were merged.
<sup>d</sup> Defined as the resolution value where the correlation between anomalous differences drops below 30 % and as calculated with program SHELXC [62].

When the six monomers of the asymmetric unit are compared to each other then two groups of molecules can be identified (group 1: chains A, C and E; group 2: chains B, D and F). Molecules within each group can be pairwise superimposed with low rmsd values ranging from 0.2 to 0.3, whereas rmsd values around 1.0 Å are obtained when comparing molecules between groups (calculated with Cα-atoms, only) (Table S4). The main divergences between the two groups occur in loop regions, namely in the loop connecting β1 to α1 and in the β2 to α2 loop.

Each dimer is formed by one molecule from group 1 and one molecule from group 2. Low rmsd values of around 0.2 Å are obtained when comparing the dimers in such a way that monomers belonging to the same group are superimposed. In contrast, when monomers are superimposed crosswise then values of around 2.6 Å are obtained. These results show that the SiiA-PD dimers are slightly asymmetric. At the same time this asymmetry is highly conserved among all three dimers present in the asymmetric unit (Table S5).

The SiiA-PD monomer structure displays a central β-sheet, consisting of 5 β-strands with mixed parallel and antiparallel strand pairing and with a β0, β1, β4, β2, β3 strand order (Fig. 2a). On one side, the β-sheet is flanked by two α-helices (α1, α2). These interconnect strands β1 and β2 (helix α1) and strands β2 and β3 (α2) (Fig. 2b). Homodimer formation is accomplished through an antiparallel edge to edge pairing of β3 from two adjacent SiiA molecules and results in the formation of a contiguous 10-stranded β-sheet (Fig. 2c). Overall, the SiiA-PD dimer displays nearly C_2_ point group symmetry with the 2-fold symmetry axis oriented perpendicular to the plane of the contiguous β-sheet. Next to β3, helix α2 also participates in the dimer interface and makes ample contacts with the homologous helix α2 from the second protomer. As shown in Fig. 2c, dimerization leads to the formation of a molecular surface that displays four α-helices and a surface which is composed of β-strands, only. Each monomer buries 780 Å^2^ of its surface in the dimer interface, and this amounts to about 12 *%* of the total monomer surface. Eleven hydrogen bonds and two salt bridges are formed between the two monomers in the dimer interface. The presence of a high number of additional hydrophobic interactions suggests that the SiiA-PD dimer constitutes a permanent dimer [28]. These results confirm earlier data where stable homodimer formation of full length SiiA was demonstrated using bacterial two hybrid assays [10]. Hence, it appears highly unlikely that transient self-association and re-dissociation events contribute to the cellular function of SiiA in *Salmonella*.

**Fig. 2.**
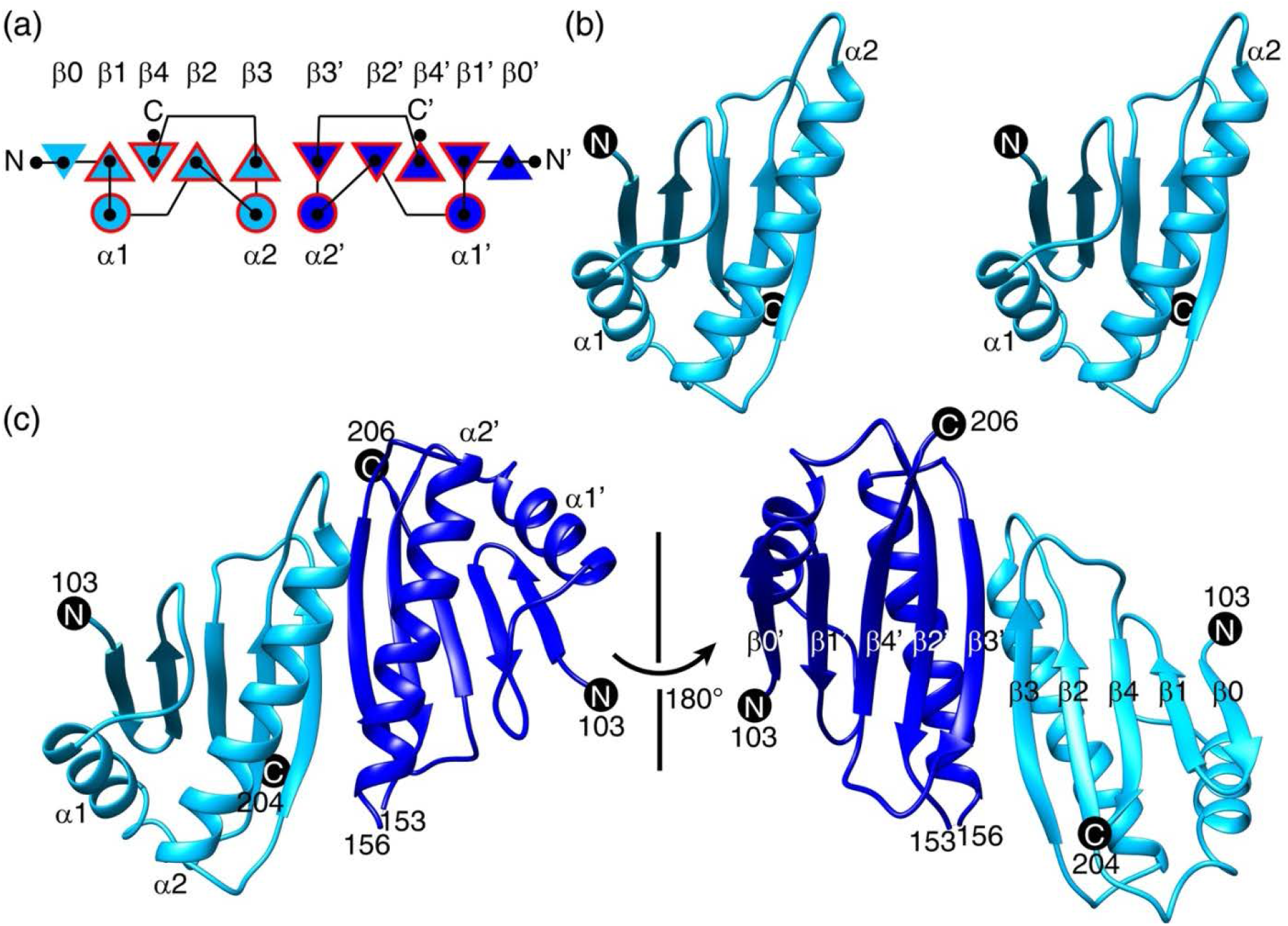
Crystal structure of SiiA-PD. (a) Topology plot of the SiiA-PD monomer (light blue) and dimer (light and dark blue). The red frame highlights the core domain fold observed in OmpA-like domains consisting of β1/α1/β2/α2/β3/β4 (see also Fig. 3). (b) Stereo image of a SiiA-PD monomer. The N-and C-termini in (b) and (c) are marked with black circles. (c) Ribbon representation of the SiiA-PD dimer. The two chains are coloured in light and dark blue.

### SiiA-PD shares structural homology to numerous PG-binding proteins

The monomer structure of SiiA-PD was used as a search template to identify homologous structures with the DALI server [29]. This search revealed that SiiA shares structural homology with numerous OmpA-like PG-binding proteins (Table 2). Several of the identified proteins were superimposed onto SiiA using the MatchMaker program in Chimera (Fig. 3, Fig. S2) [30, 31]. All of these OmpA-like domains share a common core of secondary structure elements with a β1/α1/β2/α2/β3/β4 topology. Depending on the protein, additional secondary structure elements also occur. Thus, SiiA-PD harbours an additional strand β0 at the N-terminus (Fig. 2a). OmpA from *S*. Typhimurium (PDB-ID: 5VES, [32]) and *A. baumanii* (PDB-ID: 3TD5) display an additional α3 helix present between β-strands β3 and β4 (Fig. 3). MotB from *S*. Typhimurium (PDB-ID: 2ZOV) harbors additional α0 and β0 secondary elements at the N-terminus as well as an additional α3 helix at the C-terminus (Fig. 3).

**Fig. 3.**
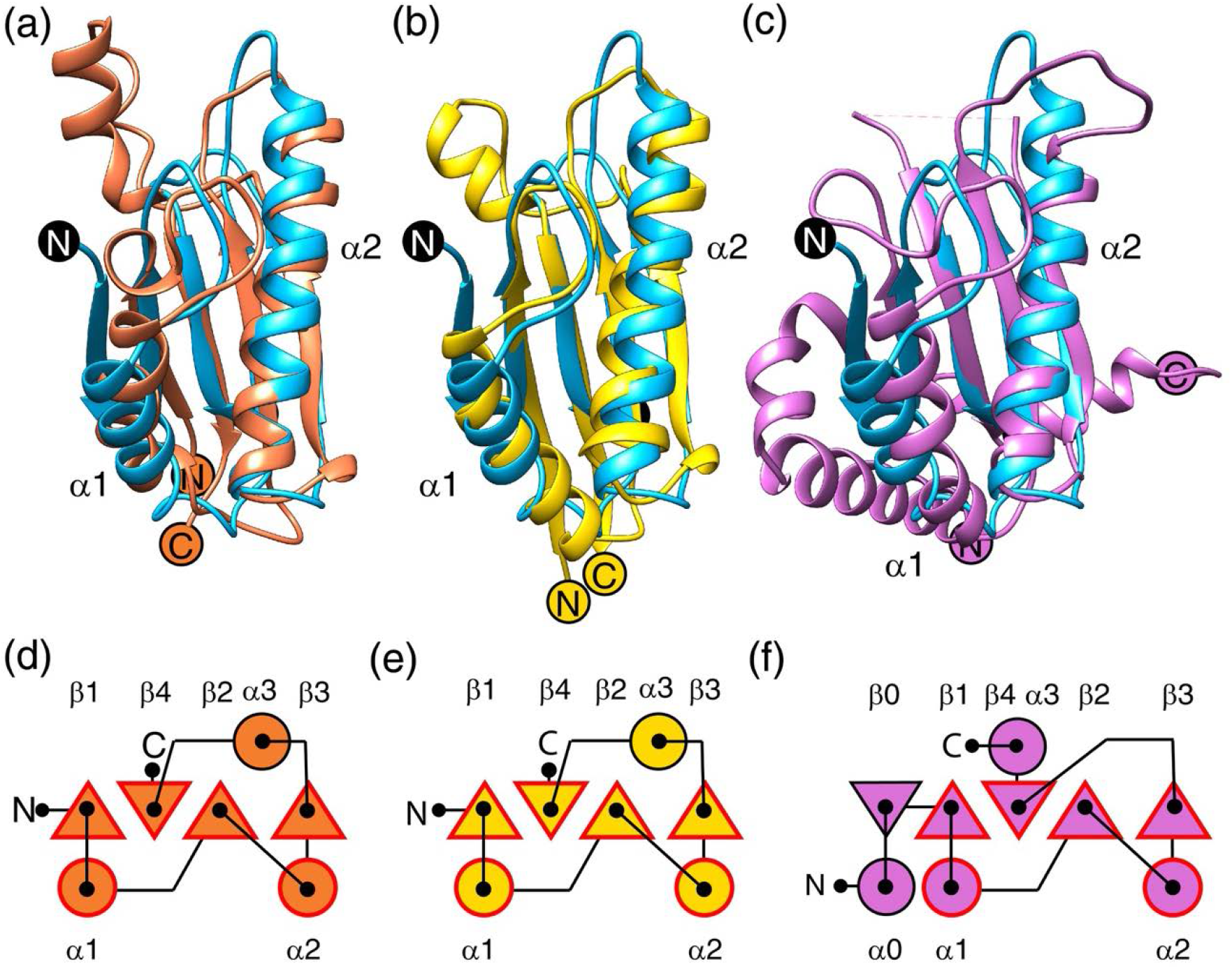
SiiA shares structural homology to OmpA-like PG-binding domains. SiiA-PD (in light blue) is structurally homologous to the PG-binding domain of (a) OmpA from *S*. Typhimurium (PDB-ID: 5VES, orange), (b) OmpA from *A. baumannii* (PDB-ID: 3TD5, yellow), and (c) MotB from *S*. Typhimurium (PDB-ID: 2ZOV, orchid), among others (Table 2, Fig. S3). N- and C-termini are marked with black circles in SiiA-PD and with coloured circles in case of the compared proteins. Helices α1 and α2 of SiiA-PD are annotated. Panels (d), (e), and (f) depict the secondary structure topology of the SiiA homologous proteins shown in panels (a), (b), and (c), respectively. All these proteins share an OmpA-like fold, consisting of a common core of β1/α1/β2/α2/β3/β3 secondary structure elements (framed in red).

**Table 2.** SiiA homologous structures.

| Protein | PDB-ID | Z-score | Cα-rmsd (Å) <sup>a</sup> | Seq. -ID (%) | Organism | Reference |
| --- | --- | --- | --- | --- | --- | --- |
| YfiB | 4zhw | 9.9 <sup>b</sup> | 2.3 (91/168) | 8 | *P. aeruginosa* | [63] |
| OmpA | 5nhx | 9.7 | 2.4 (93/344) | 8 | *K. pneumoniae* | [32] |
| MotB | 2zov | 9.4 | 2.8 (95/309) | 6 | *S. Typhimurium* | [19] |
| PAL | 5jir | 9.3 | 2.7 (97/476) | 12 | *T. pallidum* | [64] |
| TagL | 5m38 | 9.3 | 2.9 (93/576) | 5 | *E. coli* | [32] |
| PAL | 5lkw | 9.3 | 2.5 (94/170) | 11 | *B. cenocepacia* | [36] |
| PomB | 3wpw | 9.1 | 2.9 (96/315) | 7 | *V. alginolyticus* | [12] |
| PAL | 5n2c | 9.1 | 2.4 (90/170) | 11 | *B. cenocepacia* | [65] |
| OmpA | 4g88 | 9.1 | 2.4 (94/356) | 13 | *A. baumannii* | [32] |
| OmpA | 3td5 | 9.1 | 2.3 (93/356) | 13 | *A. baumannii* | [23] |
| MotB | 3s02 | 9.1 | 2.7 (95/171) | 8 | *H. pylori* | [66] |
| OmpA | 5ves | 8.9 | 2.5 (92/350) | 9 | *S. Typhimurium* | [32] |

<sup>a</sup> Numbers in parentheses indicate the number of aligned/overall residues.
<sup>b</sup> The proteins are ranked according to their Z-scores.

When comparing the dimerization modes of OmpA-like protein domains that are homologous to SiiA-PD then it becomes obvious that the dimerization modes of the periplasmic domain of MotB (MotB-PD, *e.g*. PDB-IDs: 2ZVY, 3S02), PomB (PDB-ID: 3WPW) and OmpA from *Klebsiella pneumoniae* (PDB-ID: 5NHX) are identical to that of SiiA-PD. In these proteins, dimerization is achieved by a juxtaposition of the α2 and β3 secondary structure elements from the two protomers (see above). In contrast, only monomers are observed in the crystal structures of the structurally homologous PAL protein (e.g. PDB-IDs: 5JIR, 5LKW, 5N2C). Interestingly, a different dimerization mode is observed in two crystal structures of OmpA from *A. baumannii* (*e.g*. PDB-IDs: 4G88, 3TD5). Here, dimer formation occurs *via* a juxtaposition of strand β1. However, in PDB-entry 3TD5, a second OmpA dimerization mode can be observed, too, which is similar to that of SiiA-PD. It is also notable that, among all proteins compared here, SiiA-PD shares the highest sequence identity with OmpA (13 %, Table 2). Overall, it appears that, of the different OmpA-like domain-containing proteins, all proteins proposed to be involved in ion translocation, *e.g*. MotB, PomB and SiiA, share an identical dimerization mode.

De Mot and Vanderleyden [20] analysed the sequence of various OmpA-like domains and identified a so-called PG-binding motif with the sequence TD-X_10_-LS-X_2_-RA-X_2_-V-X_3_-L. Whereas almost all of the proteins identified by the DALI server share most of the amino acids present in this motif, SiiA harbours only a very reduced PG-binding motif restricted to the sequence L-X_3_-R (Fig. 4a,b). The general motif spans from the end of strand β2 to the middle of α2 in the canonical OmpA-like domain fold (Fig. 4c). The two residues Leu163 and Arg167 from the reduced L-X3-R motif are displayed from the first half of helix α2 in SiiA (Fig. 4c). It is notable that, even though only a very low sequence homology exists between SiiA and OmpA-like domains from other proteins, *i.e*. ranging from 6 −13 %, the overall structures of these proteins remain very similar (Fig. 4d, Table 2).

**Fig. 4.**
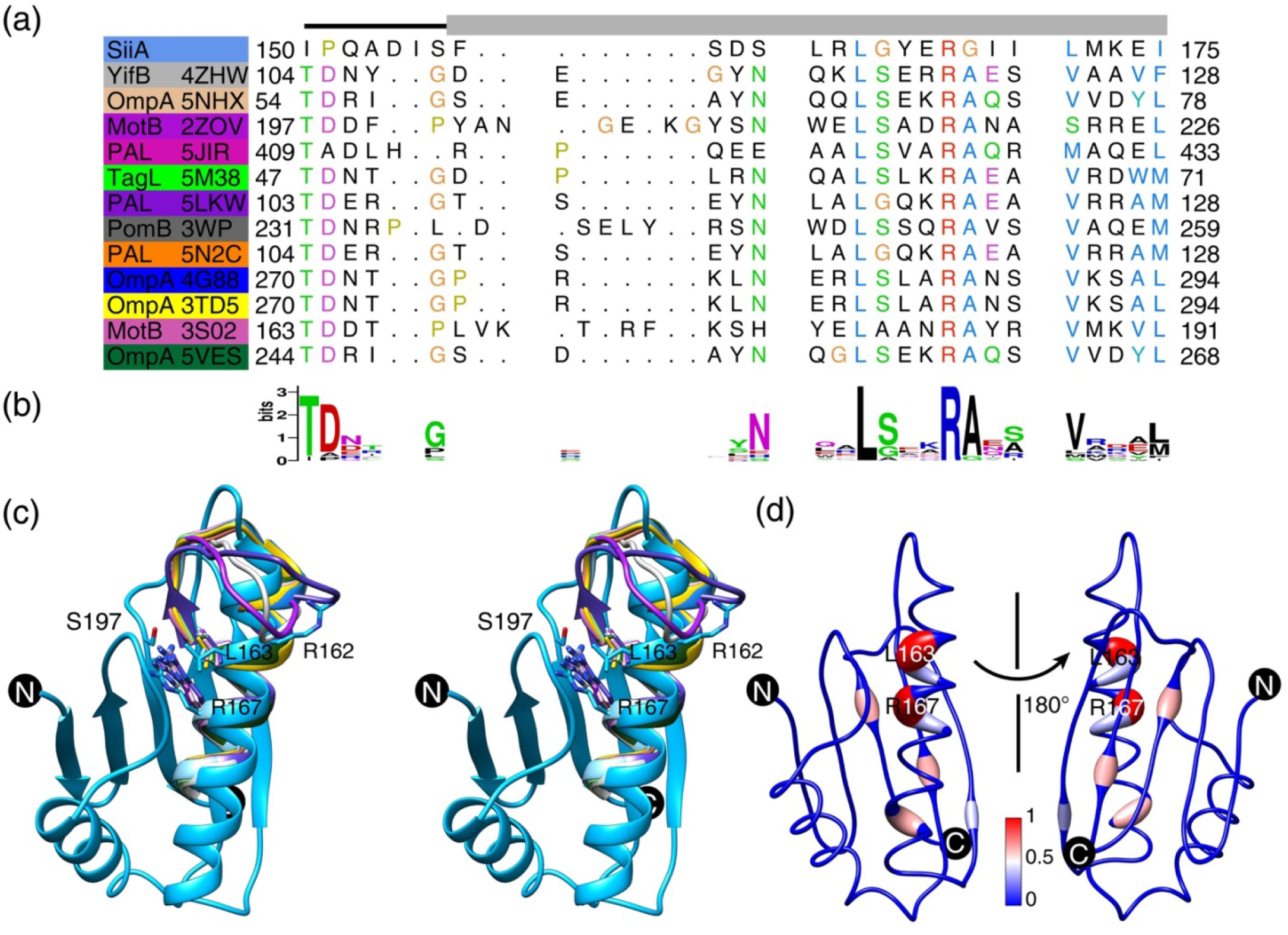
PG-binding motif of SiiA in comparison to other OmpA-like PG-binding proteins. (a) Structure-based sequence alignment of the PG-binding motif of SiiA-PD and other OmpA-like proteins. A black line (loop β2 to α2) and a grey box (helix α2) indicate the secondary structure of the aligned segment in SiiA-PD. On the left side of the alignment are reported the names and PDB-IDs of the aligned proteins (see also Table 2). (b) The PG-binding motif TDX_10_LSX_2_RAX_2_VX_3_L displayed as a sequence logo [1]. (c) Stereo image depicting the PG-binding motifs of various OmpA-like structures superimposed onto SiiA. Colour coding of the proteins as in (a). Two important residues of SiiA-PD (R162 and S197) and two selected conserved residues from the PG-binding motif (L163 and R167) are shown in a stick representation. (d) Mapping of the sequence conservation of OmpA-like PG-binding proteins onto the structure of SiiA-PD. Sequence conservation (by clustal histogram), ranging from low and high, is color-coded by a blue to red gradient and emphasized by an enlarged cartoon tube radius. The structure-based sequence alignment and the sequence conservation mapping was calculated with the full-length proteins listed in Table 2.

### SiiA-PD binds to peptidoglycan *in vitro*

SiiA-PD shares a high structural similarity with OmpA-like PG-binding domains, but, at the same time, displays a poorly conserved PG-binding motif. We used a PG pulldown assay in order to assess whether SiiA-PD actually binds to PG isolates from *S*. Typhimurium. For comparison and as a control, we also monitored binding of MotB-PD from *S*. Typhimurium to PG (residues 99 to 276 of MotB, fragment identical to that in PDB-ID: 2ZVY [19]). The binding behavior of the latter has been extensively studied before (for a recent review see [21]).

We observe that SiiA-PD is able to bind PG, albeit in a highly pH-dependent manner. SiiA binds to PG in a potassium phosphate-based buffer at a weak acidic pH, *i.e*. 5.8, whereas no binding is observed at a weak basic pH, *i.e*. 8.0. (Fig. 5, Fig. S3). This contrasts with the behaviour of MotB-PD, which binds to PG at pH 5.8 and 8.0 (Fig. 5). Binding of SiiA-PD to PG is fully reversible. When transferring the PG-bound SiiA-PD sample from pH 5.8 to 8.0 then a rapid release of PG-bound SiiA-PD into the supernatant can be observed (Fig. 5d).

**Fig. 5.**
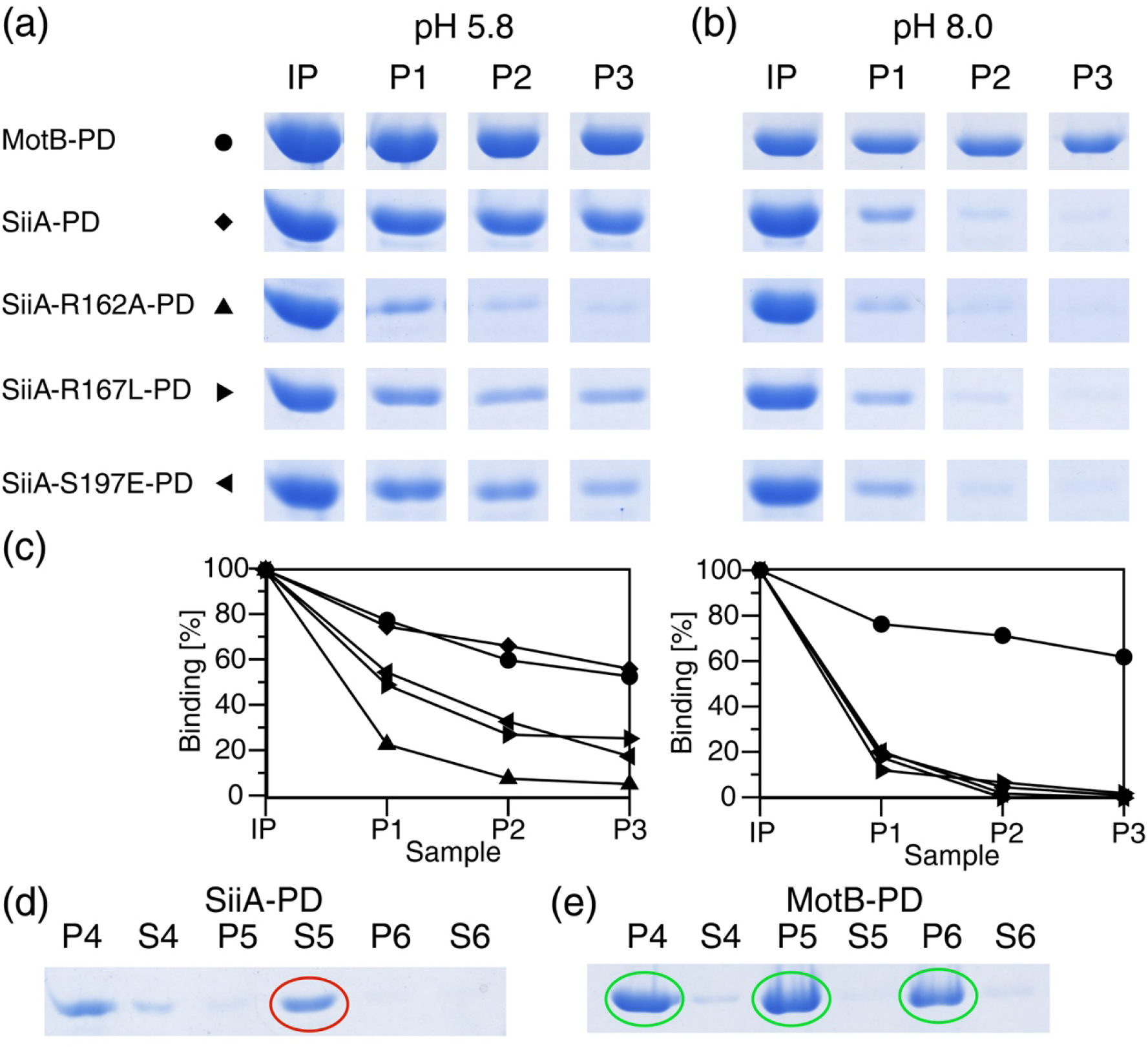
SiiA binds to PG in a pH-dependent manner. (a) and (b) PG pulldown assays of MotB-PD (dot symbol), SiiA-PD (diamond), SiiA-R162A-PD (triangle pointing upwards), SiiA-R167L-PD (triangle pointing right) and SiiA-S197E-PD (triangle pointing left) performed at pH 5.8 (a) and pH 8.0 (b). Input (IP) and pellet (P1-3) bands of the SDS-PAGE analysis of the pulldown fractions are shown. The complete SDS-PAGE gels of these assays are depicted in Fig. S3. (c) Band intensity analyses of the pulldown results. The integrated intensities of the input bands of the respective proteins were set to 100 %. The symbols used for the different proteins are identical as in (a). (d) PH-dependent dissociation of SiiA-PD from PG. (e) No pH-dependent dissociation is observed for MotB-PD. In panels (d) and (e) the P3 samples of SiiA-PD and MotB-PD from panel (a) were washed three times with a pH 8.0 buffer. P4-6 and S4-6 indicate pellet and supernatant fractions, respectively. Whereas SiiA-PD dissociates from the PG upon washing with pH 8.0 (S5, red circle), MotB-PD remains attached to PG after all washing steps (green circles).

In order to rule out that co-sedimentation of PG and SiiA-PD is caused by a pH-induced denaturation and precipitation of SiiA-PD, we investigated the stability and solubility of SiiA-PD (and MotB-PD) at pH 5.8 and 8.0 using CD spectroscopy. Over the whole pH range, both proteins show well-defined CD spectra (Fig. S4). In addition, the T_M_ values of both proteins remain unaltered at pH 5.8 and 8.0. These were measured in a CD-monitored thermal unfolding experiment and correspond to 77°C and 74°C (SiiA-PD), and 63°C and 63°C (MotB-PD) at pH 5.8 and 8.0, respectively (Fig. S4, Table S1).

These results show that the observed structural similarities between SiiA-PD and MotB-PD are paralleled by a shared function, namely the ability to bind to PG. However, the two proteins display considerably different pH-dependent affinity profiles.

### Arg162 and Arg167 participate in PG binding

An arginine residue, structurally homologous to Arg167 from the reduced PG-binding motif of SiiA, has been observed to directly participate in PG binding before. Thus, in the crystal structure of the complex formed between the PG-binding domain of OmpA from *A. baumannii* and a PG-derived pentapeptide, the guanidinium group of Arg286 of OmpA directly interacts with the carboxylate group of the *meso*-A_2_pm moiety of the bound PG-derived pentapeptide fragment (PDB-entry 3TD5, [23]). A highly similar interaction also occurs in the complex formed between the PG-binding domain of Pal and a PG precursor [24].

We produced two different mutants, namely SiiA-R167L and SiiA-S197E, in order to probe any participation of Arg167 in PG binding in SiiA. The scope of the R167L substitution was to ablate the guanidinium head group of the amino acid and, at the same time, retain the hydrophobic interactions observed between the aliphatic part of Arg167 and the SiiA protein core. The goal of the S197E substitution was to shield off the guanidinium group of Arg167 without perturbing the numerous intra-protein interactions that Arg167 is involved in. The mutant proteins display CD spectra that are indistinguishable from WT SiiA. In comparison to the latter, they show either an increased (R167L) or comparable (S197E) melting temperature. This suggests that the two point mutations do not negatively affect the structural integrity of SiiA-PD (Fig. S4c,d, Table S1). In the PG-binding assay, the two mutants share a very similar PG-binding profile. As previously observed for WT SiiA, the variants do not bind to PG at pH of 8.0. At the same time, they display a reduced PG-binding ability at pH 5.8 in comparison to WT SiiA (Fig. 5, Fig. S3). These observations strongly hint that, similar to OmpA and Pal, Arg167 represents a PG-binding determinant in SiiA.

We also tested the contribution of Arg162 to PG binding und produced the mutant variant SiiA-R162A. Arg162 is a non-conserved residue that is contained within the segment that covers the extended PG-binding motif [20]. Arg162 and Arg167 are both part of helix α2 with the former residue being located closer to the dimerization interface (Fig. 2c, Fig. 4c). Interestingly, an alanine substitution of Arg162 completely abolishes PG binding at both pH 5.8 and pH 8.0 (Fig. 5, Fig. S3). Thus, it appears that the substitution of Arg162 impairs PG binding more drastically than the substitution or shielding of Arg167. To confirm that mutation of Arg162 did not impair the structural integrity of SiiA, we investigated the mutant protein with analytical SEC and CD spectroscopy. These experiments show that dimerization of SiiA is retained in SiiA-R162A (Fig. S1) and that SiiA-R162A displays a CD spectrum and melting temperature that is similar to the WT protein (Fig. S4b, Table S1).

It is possible that Arg162 contributes indirectly to PG binding, namely through long-range electrostatic interactions. Indeed, when viewing SiiA from the side from which Arg162 is displayed then a surface patch with a striking positive electrostatic potential becomes apparent. In contrast to this, the opposite side displays an extended patch of negative electrostatic potential (Fig. 6a). If the two Arg162 residues present in the dimer are mutated *in silico* to alanine then this positive patch displays a neutral to negative potential in support of the possibility that Arg162 contributes to PG binding through electrostatic effects (Fig. 6b).

**Fig. 6.**
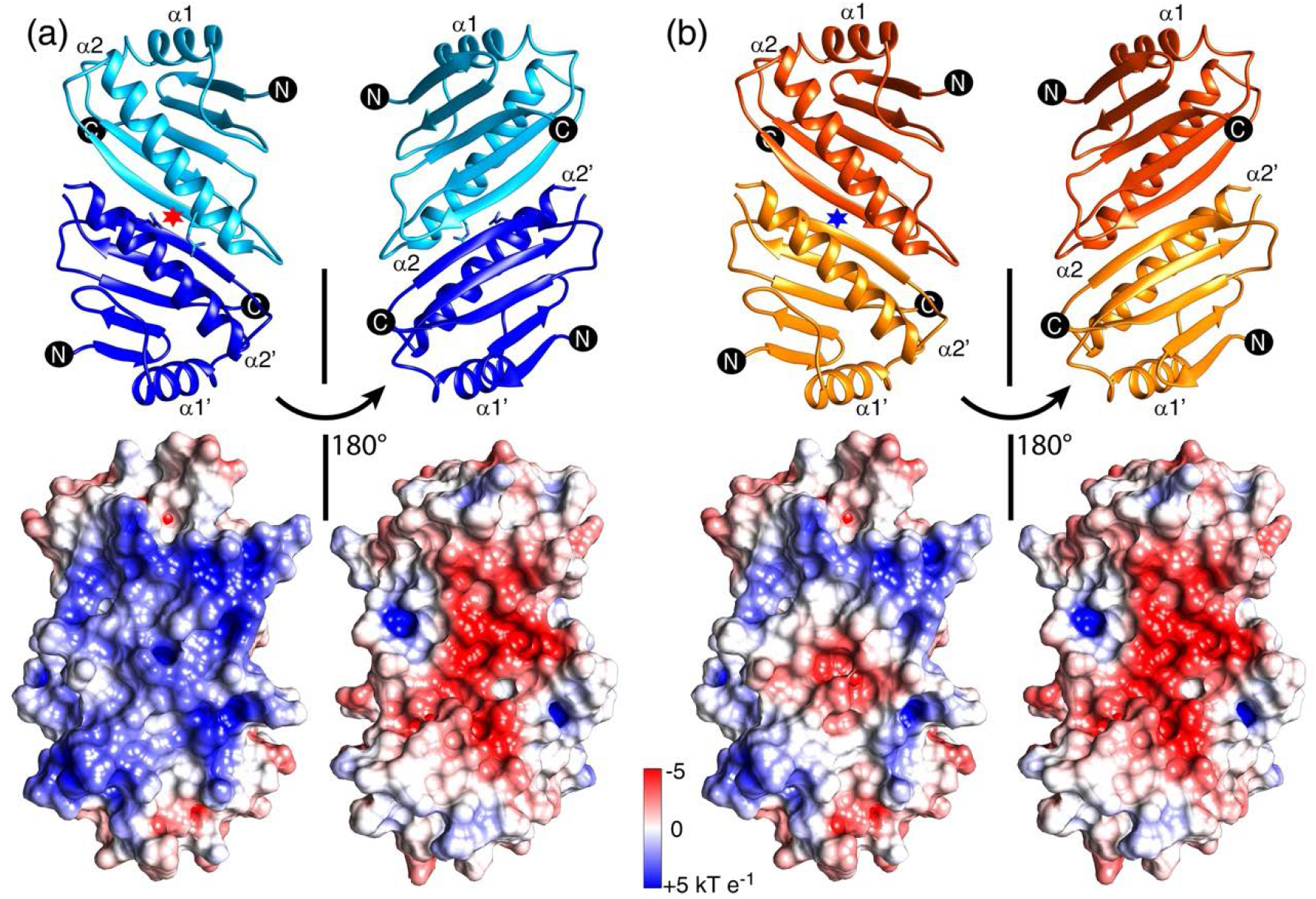
Changes in the electrostatic surface potential of SiiA-PD upon introduction of the R162A substitution. (a) The SiiA-PD dimer is displayed in light and dark blue. (b) *In silico* analysis of the R162A mutant mapped onto the SiiA-PD dimer structure shown in orange and dark yellow. In the bottom half the electrostatic surface potential is displayed from −5 to neutral to +5 kT e^-1^ with a red-white-blue color gradient. Residue R162 in panel (a) and *in silico* mutated residue A162 in (b) are shown as sticks and further highlighted by a red and blue star, respectively.

A co-isolation of either WT SiiA or mutant variant SiiA-R162A in complex with PG directly from *S*. Typhimurium further corroborates the involvement of Arg162 in PG binding. The co-isolation experiment shows that the amount of co-isolated SiiA-R162A is reduced by a factor of two in comparison to WT SiiA when analysed with an antibody directed against the HA-Tag contained in either WT SiiA or the mutant variant (Fig. 7). At the same time, antibody staining against OmpA indicates that in both co-isolated samples similar amounts of PG material are present (Table S6). Taken together, these results show that SiiA binds to PG with the help of two arginine residues. Of the two arginines, Arg 162 and Arg167, Arg162 contributes more significantly to PG binding.

**Fig. 7.**
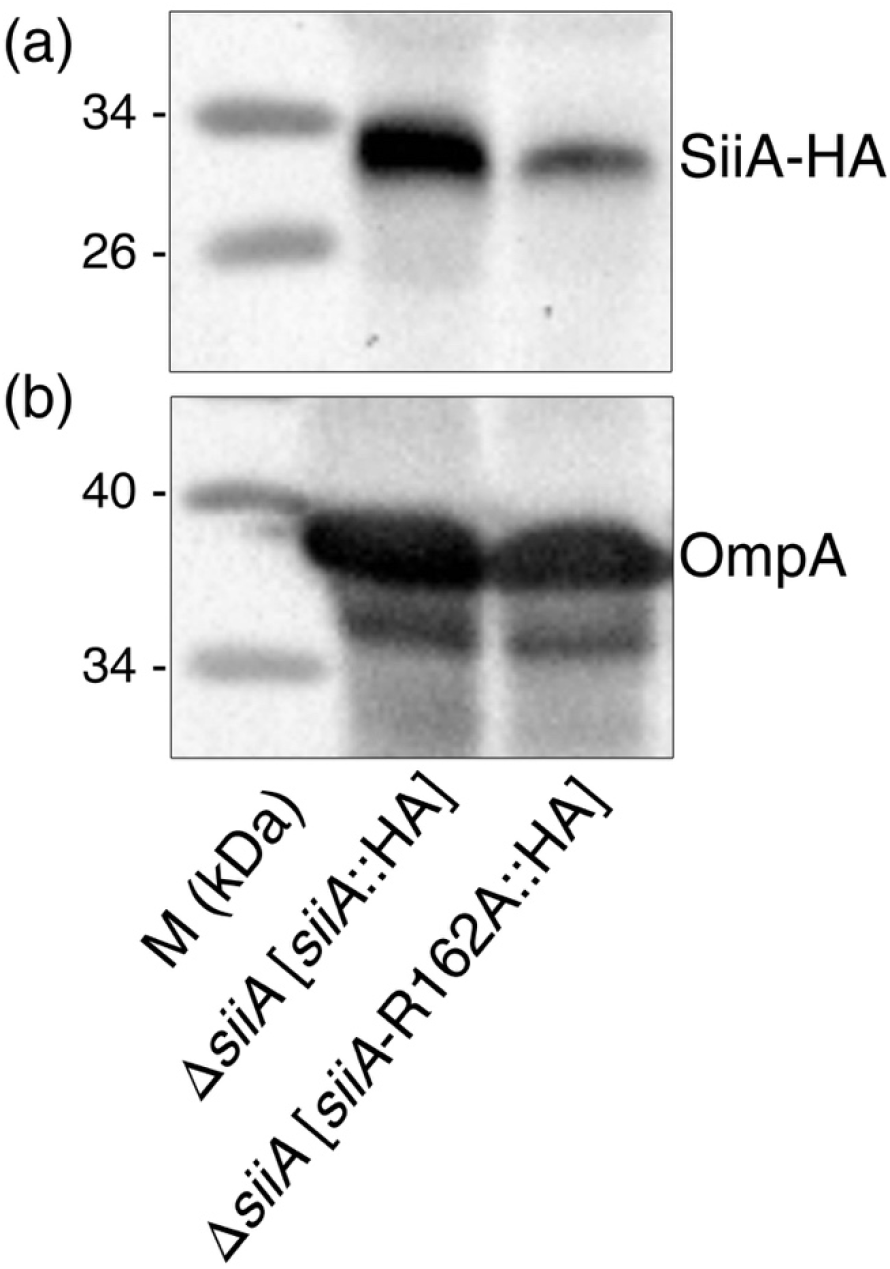
Arg162 is critical for PG binding of SiiA. *S. Typhimurium siiA* deletion strains harbouring plasmids for expression of WT *siiA*::HA or *siiA*-R162A::HA were subcultured to induce expression of genes of the SPI4-T1SS. Cells were harvested, lysed by sonication and cell wall material was isolated. PG-binding proteins were extracted by incubation with puffer containing 2 % SDS and TCA precipitated. Pellets of equal amounts of bacteria were separated by SDS-PAGE and transferred onto nitrocellulose membranes. (a) SiiA was detected by anti-HA antibody. (b) The blot was stripped and reprobed for detection of OmpA by polyclonal antiserum.

### A substitution of Arg162 reduces the *Salmonella* invasion of polarized epithelial cells

Previous studies demonstrated that the concerted actions of SiiE, the cognate T1SS and SiiAB are required to mediate adhesion to, and invasion of polarized epithelial cells [4, 6]. Mutant strains deficient in *siiA* are highly attenuated in invasion of polarized epithelial cells [10]. We therefore set out to investigate if the PG-binding domain of SiiA is also important for the overall function of the SPI4-encoded T1SS. First, a mutational analysis was performed by C-terminal truncations or internal deletions to various extents of the PG-binding domain. We observed that the cellular amounts of the resulting mutant forms of SiiA were highly reduced, thus impeding further functional analyses (data not shown).

We investigated next whether a substitution of individual amino acids affects invasion of polarized epithelial cells. In order to identify candidate residues, in addition to R162A, R167L and S197E, we superimposed the structure of SiiA onto the structure of OmpA in complex with a PG fragment (PDB-entry 3TD5, [23]) and considered all SiiA residues that were located within 5 Å of the OmpA-bound ligand as potential substitution candidates.

We performed site-directed mutagenesis of *siiA* and generated R120A, R159A, R162A, L163A, R167L and S197E variant alleles. The synthesis of WT and mutant forms of SiiA was analysed in total cell lysates of *S*. Typhimurium strains. We observed that, while the cellular amounts of SiiA mutants D159A, L163A and R167L were highly reduced, the amounts of SiiA variants R120A, R162A and S197E were comparable to the levels of WT SiiA (Fig. 8a).

**Fig. 8.**
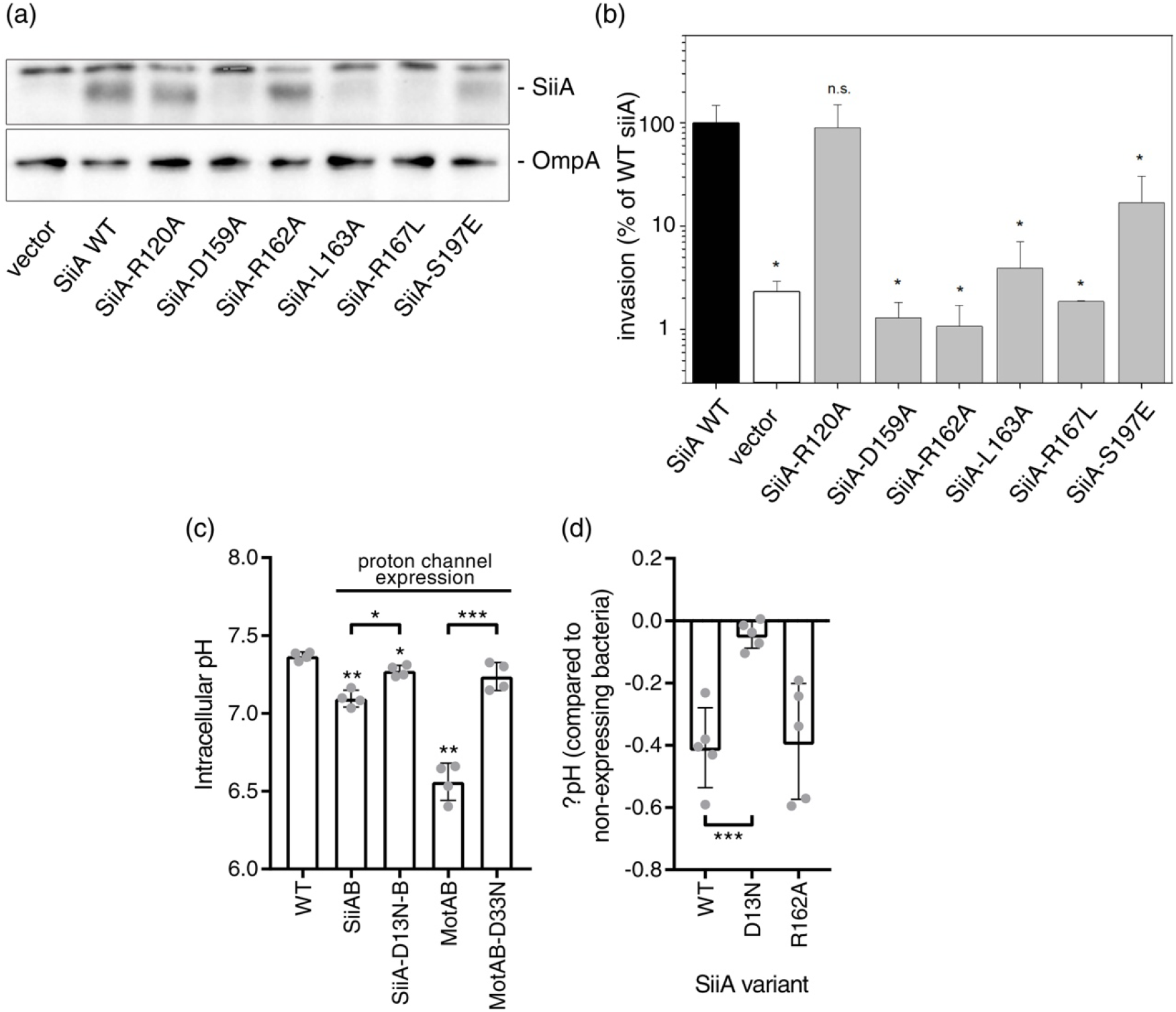
Mutational analysis of SiiA *in vivo*. (a) Effect of *siiA* site-directed mutagenesis (SDM) on cellular amounts of SiiA. *S*. Typhimurium MvP771 harbouring the empty vector or various plasmids for the expression of *siiA* WT or various mutant alleles were subcultured for LB broth for 3.5 h. Equal amounts of bacterial cells as adjusted by OD600 were harvested and by SDS-PAGE sample buffer. Protein was separated by SDS-PAGE on 12 % gels and transferred onto nitrocellulose membranes. SiiA was detected using antibody against the HA tag. As loading control, blots were stripped and developed using an antibody against DnaK. (b) Effect of SDM of *siiA* on SPI4-dependent invasion of polarized epithelial cells. The Δ*siiA* strain MvP771 harbouring plasmids for the expression of WT *siiA*, various mutant alleles, or the empty vector were subcultured for 3.5 h to induce expression of genes for adhesion to, and invasion of polarized epithelial cells. The canine kidney epithelial cell line MDCK was grown as polarized monolayer and infected with *Salmonella* at a multiplicity of infection (MOI) of 5. After incubation for 25 min, non-internalized *Salmonella* were removed by washing and remaining extracellular bacteria were killed by incubation in medium containing 100 μg/ml Gentamicin for 1 h. Cell were washed again and lysed by addition of PBS containing 0.5 % desoxycholate. Lysates for plates onto Mueller-Hinton agar to determine the number of colony forming units. Invasion is expressed as percentage of internalized inoculum. Means and standard deviations of triplicate samples are shown and the data are representative for three independent experiments. (c) Intracellular pH was detected using ratiometric fluorescence measurement of R-pHluorin-M153R in *S*. Typhimurium NCTC 12023 without (WT) or with co-expression of SiiAB, SiiA-D13N-SiiB, MotAB or MotA-MotB-D33N as indicated. Data of four independent experiments are shown and statistical significance was calculated against WT or between indicated data sets using paired, two-tailed Students *t*-test and was defined as * for *p* < 0.05, ** for *p* < 0.01 or *** for *p* < 0.001. (d) Change in intracellular pH after expression of SiiAB complexes with WT SiiA, SiiA-D13N or SiiA-R162A compared to bacteria without induction of expression. Data of five independent experiments are shown and statistical significance was calculated using unpaired, two-tailed Students *t*-test defined as *** for *p* < 0.001.

We compared phenotypes of a *siiA* deletion strains harbouring plasmids for the production of WT SiiA and SiiA mutants and studied the effect of aa exchanges in SiiA on invasion of polarized cells (Fig. 8b). SiiA-R120A and SiiA-S197E restored invasion similar to WT SiiA. SiiA-D159A, SiiA-R162A, SiiA-L163A and SiiE-R167L did not restore invasion and entry levels were comparable to a strain harbouring the empty vector. However, since the cellular amounts of SiiA-D159A, SiiA-L163A and SiiE-R167L were highly reduced, it is not possible to estimate the contribution of the mutated residues to SiiA function.

In contrast, SiiA-R120A, SiiA-R162A, and SiiA-S197E levels were not decreased compared to WT SiiA. Of these three mutants, only the mutant R162A showed reduced invasion. Thus, we conclude that residue Arg162 is critical for the function of SiiA, while amino acid exchanges R120A and S197E are compatible with SiiA function. It is interesting to note that substitution of Arg162 leads to both an abolishment of PG binding and a reduction of the invasion efficacy, whereas substitution of S197E causes only a reduction of the PG-binding affinity.

### Transmembrane substitutions alter the proton-conducting activity of SiiAB

Encouraged by the anticipated structural similarities between SiiAB and MotAB and ExbBD, we previously suggested that SiiAB resembles a proton-conducting channel in the IM [10]. In order to test this proposed function of SiiAB, we established an assay that allows assessing the impact of SiiAB onto the intracellular pH of living bacteria, as previously demonstrated for MotAB [33]. For that, a pH-sensing GFP variant, namely R-pHluorin-M153R, was co-produced with either SiiAB or MotAB [34]. Co-production of either complexes led to a significantly more acidic cytosolic pH, which argues for the occurrence of a functional proton influx through both proton channels (Fig. 8c).

Consistent with the proposed crucial role of a transmembrane aspartate in proton conductance [10, 11], the acidification of the cytoplasm was strongly impeded when either the mutant complexes MotA-MotB-D33N or SiiA-D13N-SiiB were expressed (Fig. 8c). At the same time, WT and mutant SiiA proteins showed equal expression levels as detected by Western blot (Fig. S5).

In a similar set of experiments, we measured the intracellular pH of *Salmonella* expressing SiiAB-complexes containing the R162A substitution in the PG-binding domain of SiiA. Expression of SiiA-R162A-SiiB was controlled again *via* Western blot (Fig. S5). Expression of the mutant induced a cytosolic acidification comparable to that observed for WT SiiAB (Fig. 8d). Thus, no distinction can be made between the behaviour of WT SiiA and SiiA-R162A with regard to cytosolic acidification.

## Discussion

In the present study, we investigated the atomic structure and function of SiiA, a protein that forms a complex with SiiB in the IM [10]. The SiiAB complex shares a number of properties with MotAB as well as with other members of the family of heteromeric ion-conducting channels, such as PomAB, ExbBD and TolQR. These latter complexes, by means of ion translocation across the IM, generate mechanical work that translates into the rotation of the flagellum or the uptake of nutrients at the OM [12, 17, 19, 35]. The question arises whether any structural similarities between SiiAB and for example MotAB are paralleled by similarities in the regulatory mechanisms that these proteins contribute to. If so, then available knowledge on MotAB could help to better understand how SiiAB contributes to SPI4-mediated adhesion of *Salmonella* [10].

Here, we experimentally showed that indeed the architecture of SiiA closely resembles that of MotB. SiiA consists of a single transmembrane domain followed by an intrinsically disordered region and a stably-folded domain referred to as SiiA-PD. Prior work demonstrated that residue Asp13 in the transmembrane region of SiiA is essential for SiiA function. Exchange of Asp13 in SiiA resulted in a highly attenuated invasion of polarized epithelial cells. Here, we now showed that the complex SiiAB is indeed involved in proton translocation across the IM. However, at the pH value tested, *i.e*. pH 7.4, SiiAB was not able to reduce the cytosolic pH to the same extent as MotAB (Fig. 8c) [33]. This could be caused by the reduced PG-binding affinity of SiiA at neutral pH. This would also explain why mutant SiiA-R162A, for which we observed an impeded PG binding, behaves quite similar in this assay. Clearly, proton influx at a pH where SiiA binds to PG remains to be analysed. This notwithstanding, our experiments show that Asp13 in SiiA as well as Asp33 in MotB directly imped proton influx. Therefore, mutation of Asp13 in SiiA affects both proton translocation and, as previously shown, invasion of polarized epithelial cells [10]. These results strongly hint that proton translocation through SiiAB and hence energy derived from the PMF directly contribute to SPI4-mediated adhesion of *Salmonella*.

We solved the crystal structure of the stably-folded periplasmic domain of SiiA, *i.e*. SiiA-PD. The structure of SiiA-PD resembles the structures of PG-binding domains from proteins present in either the IM or OM of Gram-negative bacteria. SiiA-PD forms a homodimer, and an identical dimerization mode is observed in MotB and PomB. In contrast, the dimerization mode of the PG-binding domain of OmpA, a protein located in the OM, is remarkably different. The OmpA-like domain of PAL, on the other hand, is monomeric in solution and in the crystal structure [36]. Thus, it appears that PG-binding domains of proteins involved in ion translocation across the IM, *i.e*. MotB, PomB and SiiA, share the same dimerization mode, whereas this is not the case for proteins with other functions.

SiiA harbours a reduced PG-binding motif. Only two residues, namely Leu163 and Arg167, are strictly conserved between the motif in SiiA-PD and the motif in other OmpA-like PG-binding domains (Fig. 4a). Here, we now showed for the first time that SiiA binds indeed to PG *in vitro* and that mutation or the shielding-off of Arg167 leads to a reduction of its PG-binding affinity. Moreover, mutation of additional residue Arg162, located in close proximity to Arg167, completely abolished PG binding *in vitro*. SiiA bound to PG with higher affinity at low pH (pH 5.8), whereas MotB that we used as a control protein bound PG at low and high pH (pH 5.8 and 8.0). We also observed that increasing the salt concentration in the SiiA buffer negatively affected PG binding of SiiA (data not shown). It is difficult to anticipate how this *in vitro* observation translates into the number of PG-bound *versus* unbound SiiA molecules *in vivo*. Free diffusion of SiiA is restricted by both the attachment of SiiA to the membrane and its confinement to the periplasmic space. As a result, high local apparent concentrations prevail. Hence, the fraction of PG-bound SiiA molecules could by high despite SiiA displaying only a moderate PG-binding affinity under physiological conditions [37].

Our results show that pH modulates the PG-binding affinity of SiiA, a property so far not observed for other PG-binding proteins. Notably, the binding affinity increases at low pH (Fig. 5). Taking the oral infection route, *Salmonella* survive the highly acidic pH in the stomach and are exposed to a rising pH gradient within the gut lumen reaching neutral levels in the ileum and decreasing again to mildly acidic levels in the colon [38]. So *Salmonella* are exposed to a low pH environment *in vivo* before it gets neutralized by bicarbonate-containing mucus overlaying the enterocytes [39]. It remains to be determined how the observed pH dependency of SiiA binding to PG translates into the detailed mechanism of the infection process *in vivo* since to the present day infection experiments on polarized MDCK cells are usually performed at neutral pH [4].

Our experiments suggest that the segment that links the transmembrane region of SiiA to the stably folded SiiA-PD is intrinsically disordered. Bioinformatics analyses, limited proteolysis experiments and difference CD spectra fail to reveal the presence of any secondary structure elements in this segment. This appears to be different for other members of this family. In MotB, PomB and TolR, this segment is proposed to play an important role in regulating both the anchoring of the PD domain to the PG layer and in controlling the translocation of ions across the translocation pore [12, 17, 19]. In the inactive form of MotB, a so-called plug helix, contained in the segment that interlinks the transmembrane region of MotB to MotB-PD, is proposed to obstruct the proton-pore and inhibit proton translocation. Once the MotB-binding partner MotA interacts with the flagellum protein FliG, a structural rearrangement occurs that opens both the channel and induces a conformational change in the linker segment allowing for the attachment of MotB-PD to the PG layer. Overall, highly similar mechanisms have been proposed for how the linker segment controls the PG layer attachment in MotB, PomB and TolR [12, 17, 19, 40]. In these proteins, the function of the linker segment resembles that of a train pantograph. The latter either sits flat on the train roof (PG-domain remaining attached to the IM) or extends and connects the train to the overhead electric line (PG-domain attached to the PG layer).

Our present experiments fail to reveal that the segment that interconnects the transmembrane region of SiiA to the SiiA-PD is able to adopt different conformations. Moreover, MotB-SiiA chimeric proteins, in which the MotB PEM has been substituted against corresponding segments from SiiA, are not able to rescue flagellum function in the Δ*motAB* mutant (data not shown). Nevertheless, it appears tempting to speculate that the acidification of the periplasmic space of *Salmonella* in the gut could provide for a regulatory trigger signal. As a result of the acidification, SiiA-PD attaches to the PG layer, and the 60 residue-long disordered linker segment is forced into an extended conformation that triggers the opening of the proton pore. If this holds true then also in SiiA the linker segment would assume a regulatory role. However, the primary trigger signal for the opening of the SiiAB pore would be the acidification of the periplasm, a mechanism so far not observed in MotB, PomB and TolR (Fig. 9).

**Fig. 9.**
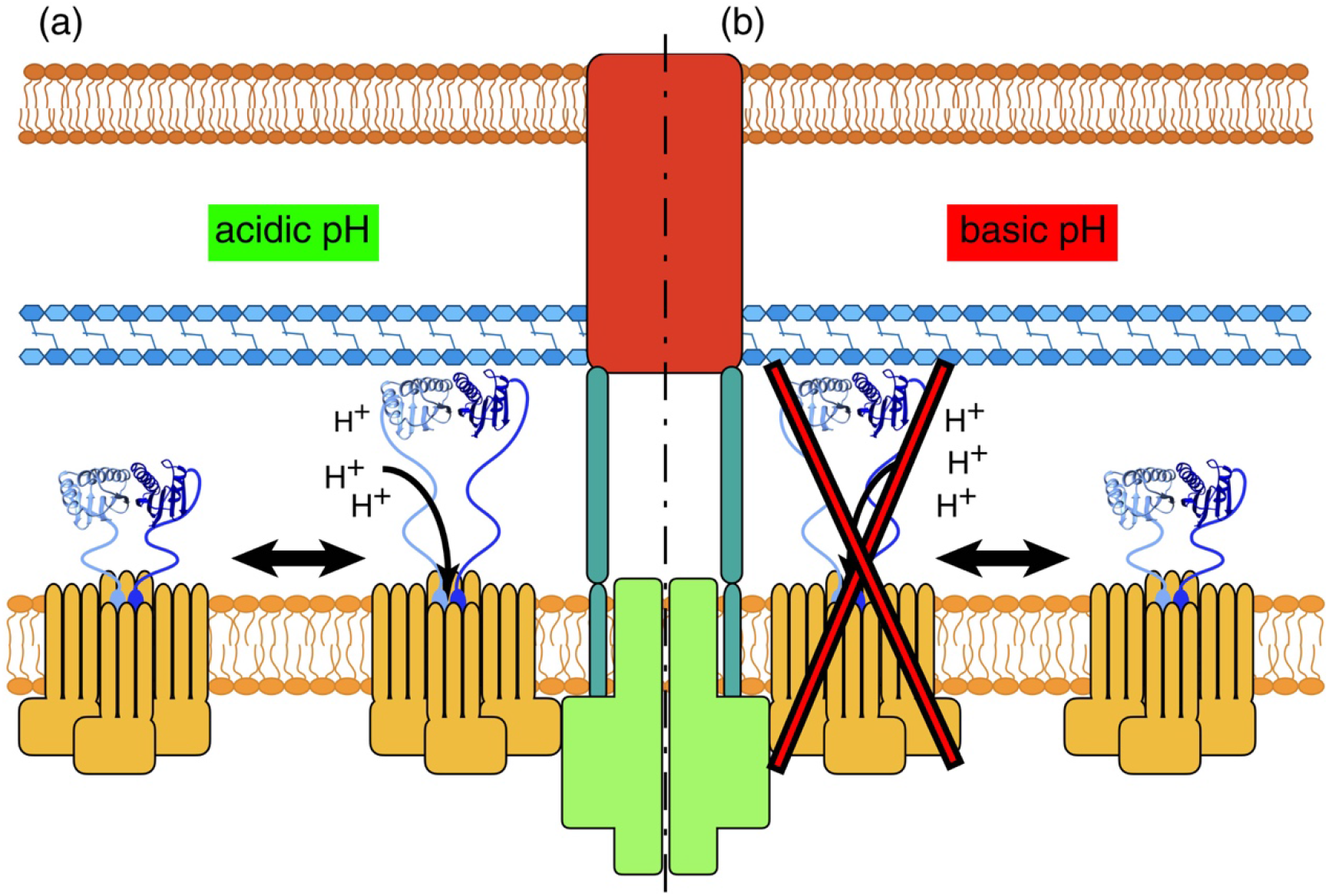
Model of the SiiAB-complex and its integration into the SPI4-T1SS. (a) Upon acidification of the periplasmic space SiiA-PD attaches to the PG-layer and to SiiF and the proton influx is enabled. (b) At a basic pH, SiiA-PD is not able to bind to PG and the proton influx is reduced. At present it is still not known how proton influx or possibly any work that is generated by the influx of protons contribute to the function of the T1SS. SiiA and SiiB are shown in blue and yellow, respectively. The canonical components of the SPI4-encoded T1SS are shown in red (SiiC), light green (SiiF) and cyan (SiiD).

Presently, it still remains unclear how exactly SiiAB contributes to the function of the SPI4-encoded T1SS and ultimately invasion of epithelial cells by *Salmonella*. Here and in past experiments, it is shown that the participation of SiiA in proton translocation and the ability of SiiA to attach its PD to the PG layer contribute to proper T1SS function. Moreover, the structural and functional similarities observed between SiiAB and MotAB strongly suggest that SiiAB is able to ‘tap’ into the PMF. It remains to be shown whether this translates into mechanical work, and, if so, how any SiiAB-produced mechanical work contributes to *Salmonella* infectivity.

## Material and Methods

### Protein production

For the production of ppr-SiiA, the *siia* gene was cloned into a pET15b vector (Novagen, Darmstadt, Germany). The translated gene product consists of an N-terminal hexa-histidine tag (MGSSHHHHHH) followed by a thrombin cleavage site (SSGLVPRGSHM) and residues 40 to 210 of SiiA from *S*. Typhimurium (Uniprot entry H9L4G5, [41]). The SiiA-R162A mutant was generated using a 2-stage mutagenesis protocol [42]. Two separate 50 μL PCR mixtures, containing either the forward or the reverse PCR primer listed, were prepared and 7 PCR cycles performed (Table S7). Subsequently, 25 μL of each mixture were combined, additional Q5 polymerase (NEB, Frankfurt, Germany) added and 17 additional PCR cycles performed. DpnI (NEB) was added, and the mixture incubated at 37°C for 1 h prior to the transformation into XL10Gold cells (NEB) (Table S8). The correctness of all plasmid constructs generated in this study was confirmed by DNA sequencing.

Additional mutant variants were generated with the oligonucleotides listed in Table S7 and using a protocol that adhered to the manufacturer’s instructions of the Q5 SDM kit (NEB). The protocol contained several modifications: the kinase-ligase-DpnI reaction was performed by mixing 10, 400 and 20 units of T4-polynucletotide kinase, T4-ligase and DpnI (all NEB), respectively. The reaction was incubated for 2 h at room temperature (RT). 5 μL of this reaction mixture were transformed into XL10Gold cells (NEB).

WT ppr-SiiA protein and mutant variants were produced using freshly transformed BL21(DE3) (Novagen) cells. The transformed cells were incubated in LB medium (Carl Roth, Karlsruhe, Germany) supplemented with 100 μg/mL ampicillin. The cells were harvested by centrifugation, resuspended in a sodium phosphate/imidazole buffer (20 mM sodium phosphate pH 7.5, 30 mM imidazole, 500 mM NaCl, EDTA-free protease inhibitor cocktail (Roche, Mannheim, Germany)) and lysed by sonication. Cell debris was removed by centrifugation at 95,000 x g and 4°C for 1 h. The proteins were purified by affinity chromatography *via* the hexa-histidine tag (elution buffer: 20 mM sodium phosphate pH 7.5, 500 mM imidazole, 500 mM NaCl) and dialysed into a 20 mM Tris-HCl pH 7.5 buffer. The hexa-histidine tag was removed overnight with thrombin (5 U thrombin (Sigma-Aldrich, Darmstadt, Germany) per mg SiiA protein) resulting in a fragment covering the periplasmic part of SiiA. The cleavage reaction was stopped by adding 5 mM AEBSF, and the protein was further purified by size exclusion chromatography using a Superdex 75 16 600 column (GE Healthcare, Freiburg, Germany) and a 20 mM Tris-HCl pH 7.5 buffer.

The production of seleno-methionine (SeMet)-labelled ppr-SiiA was accomplished using the protocol of Studier *et al*. [43]. The transformed BL21(DE3) cells were grown overnight in PAG media supplemented with 100 μg/mL ampicillin [43]. This preculture was then diluted into the auto-inducing PSAM 5052 medium and supplemented with 100 μg/mL carbenicillin [43]. The cells were grown for 3.5 days at 20°C before harvesting by centrifugation. SeMet-labelled SiiA protein isolation and purification was carried out as described above, except that the buffer used for size exclusion chromatography consisted of 20 mM Tris-HCl pH 7.5, 10 mM EDTA and 150 mM NaCl. The protein was stored at a concentration of 1.0 mg/mL at −80°C for later usage.

In order to have a positive control protein sample in the PG-binding assay (see below), the PG-binding domain of *Salmonella* MotB (aa 99-276, Uniprot entry P55892, [41]) was produced using a similar gene construct as for SiiA. MotB was cloned into a pET15b vector (Novagen) resulting in a construct that contains an N-terminal His-tag and a thrombin cleavage site. BL21(DE3) cells were transformed with the plasmids, and the transformed cells incubated in TB medium (Carl Roth) supplemented with 100 μg/mL ampicillin. The cells were harvested by centrifugation, resuspended in a Tris/imidazole buffer (20 mM Tris-HCl, 20 mM imidazole, pH 7.4, 500 mM NaCl, EDTA-free protease inhibitor cocktail) and lysed by sonication. Cell debris was removed by centrifugation at 108,000 x g and 4°C for 1 h. MotB was purified by affinity chromatography *via* the hexa-histidine tag (elution buffer: 20 mM Tris-HCl, 500 mM Imidazole pH 7.4, 500 mM NaCl) and subsequently dialyzed against 20 mM Tris-HCl pH 8.0, 100 mM NaCl buffer. MotB was further purified by size exclusion chromatography (Superdex 75 16 600 column) using a 20 mM Tris-HCl pH 8.0, 100 mM NaCl buffer.

### Proteolysis with thermolysin

In order to investigate the domain structure of ppr-SiiA and to identify alternative SiiA fragments for crystallization trials, we incubated a purified ppr-SiiA sample over a period of 2 days at 20°C with limited amounts of thermolysin (Hampton Research, Aliso Viejo, USA; 1 μg thermolysin per mg SiiA). The formation of proteolytic degradation products was monitored by retrieving aliquots at fixed time intervals and analysing the aliquots with SDS-PAGE.

For preparative production of WT and mutant SiiA-PD (either SeMet-labelled or not), thermolysin was added to ppr-SiiA (1 μg thermolysin per mg SiiA) after the initial His-tag affinity chromatography step in a 20 mM Tris-HCl pH 7.5 buffer. The thermolysin-SiiA mixture was incubated for 1.5 hour on ice. Proteolysis was stopped by adding 10 mM EDTA. The resulting protein fragment was purified using a DEAE-sepharose column (GE Healthcare). Elution from the column was accomplished with a linear gradient ranging from 0 to 500 mM NaCl in a buffer containing 20 mM Tris-HCl pH 7.4 and 10 mM EDTA. The final purification step consisted of a size exclusion chromatography (Superdex 75 16 600 column, GE Healthcare) and using the same buffer as above but without NaCl. SiiA-PD was then concentrated to 20 mg/mL and stored at −80°C for later usage.

### CD measurements

The secondary structure composition of the different protein variants was analyzed *via* CD spectroscopy, using a Jasco J-815 CD spectropolarimeter (Jasco, Gross-Umstadt, Germany). Prior to the recording of the CD signal, the protein variants were dialyzed against buffers consisting of 10 mM potassium phosphate with different pH values, namely pH 5.8, 7.4 and 8.0. All spectra were recorded in a 0.1 cm cuvette with protein concentrations of 5 μM protein. The absorption range was scanned from 185 to 260 nm using a step size of 0.1 nm, a data integration time of 1s, a bandwidth of 1 nm and a scanning speed of 20 nm/min. For further analysis of the CD signal, the mean residue ellipticity was calculated [44]. Thermal denaturation analysis was performed in a 1 cm cuvette with a protein concentration of 0.7 μM. The CD signal was recorded at a single wavelength, namely at 222 nm, and the temperature increased from 20 to 100°C at a rate of 1°C per min.

### Protein crystallization and crystal structure determination

SeMet-labelled SiiA-PD was crystallized using the hanging drop method [45]. The reservoir solution consisted of 200 mM sodium phosphate dibasic pH 8.8 ± 0.1 and 20 % PEG 3350. The protein droplet was formed by mixing 2 μL of protein solution (17 mg/mL SeMet-labelled SiiA-PD in 20 mM Tris-HCl pH 7.5, 1 mM EDTA and 15 mM NaCl) with 1 μL reservoir solution and 2 μL of perfluoropolyether cryo-oil (Hampton research). The crystals were harvested, cryo-protected with additional cryo-oil and flash frozen in liquid nitrogen prior to any diffraction data collection.

Diffraction data were collected on beamline BL14.1 operated by the Helmholtz-Zentrum Berlin (HZB) and BL14.2 operated by the Joint Berlin MX-Laboratory at the BESSY II electron storage ring (Berlin-Adlershof, Germany) [46]. The structure of SiiA-PD was solved using the multi-wavelength anomalous dispersion (MAD) method [45]. A complete MAD dataset was collected from a single triclinic SeMet-labelled SiiA-PD crystal at beamline BL14.2, and the data processed with XDS and XDSAPP [47, 48]. The values of the triclinic unit cell dimensions (*a, b, c* and *α, β, γ* equal to 58.3, 58.4, 65.8 (Å) and 93.9, 94.0, 119.0 (°), respectively) are very close to values that would allow for an alternative crystal lattice assignment, namely a monoclinic C-centred Bravais lattice with dimensions 59.2, 100.6, 65.8 (Å) and 90.0, 97.8, 90.0 (°). However, an analysis of the structure factor amplitudes of the native dataset with programs XDS and POINTLESS revealed unambiguously the absence of a two-fold rotational symmetry (R_meas_ ~ 11 % in Laue group −1 *versus* R_meas_ ~ 37 % in group 2/m) and confirmed the triclinic space group (Table 1) [47, 49]. Data were scaled with program XSCALE, and the localization of the selenium sites and partial model building accomplished with program AUTOSOL of the PHENIX software suite [50, 51]. Model building was completed, and the structure refined to convergence in an iterative process, consisting of steps of manual inspection with program COOT, automated model rebuilding and refinement, using program AUTOBUILD and single model refinement steps with program PHENIX.REFINE [52–54]. The triclinic crystals contain six protein molecules grouped into three dimers in the asymmetric unit. Data collection and refinement statistics are summarized in Table 1.

### Bioinformatics analyses

A monomeric chain of SiiA was used with the DALI server to identify protein structures that are homologous to SiiA [29]. The identified structures were ranked based on Z-scores. Crystal structures with a Z-score > 8.9 were chosen for a structure-based sequence alignment. The structures were aligned with the MATCHMAKER program in CHIMERA [30, 31]. The sequence conservation mapping was performed with the full-length proteins and the MULTIALIGN VIEWER in CHIMERA [30, 31].

The electrostatic surface representation was generated *via* the APBS-PDB2PQR-server version 2.1.1 with default settings (http://nbcr-222.ucsd.edu/pdb2pqr_2.1.1/) [55]. The PDB2PQR settings included the use of the PARSE-force field [56], the treatment of the N- and C-terminus as neutral and a pH of 7.0.

Rmsd values between monomers and dimers were calculated with LSQKAP [57]. Changes in solvent-accessible surface areas were calculated with the PISA-server [58]. All structure illustrations were drawn with program chimera [30].

### Peptidoglycan (PG) isolation

The isolation of PG from *S*. Typhimurium was performed according to a protocol adapted from Glauner (1988) [59]. In short, bacteria were grown in LB medium (Carl Roth) to mid-exponential log phase (OD_600_ of 0.6-0.8). The cells were chilled rapidly in an ice bath and harvested via centrifugation. The cell pellet was resuspended in MQ-water and added dropwise to a boiling SDS-solution (8 %). The volume was adjusted with MQ to 4 % SDS and further boiled for 30 min. From now on, the solution was kept at RT. The next day, the solution was centrifuged at 108,000 x g at RT for 45 min. The pellet was washed with MQ-water five times in order to remove any residual SDS. The PG pellet was suspended in 50 mM Tris-HCl pH 7.0 and subsequently treated with 100 μg mL^-1^ α-amylase (Sigma-Aldrich) for 2 h at 37°C. The solution was then supplemented with 20 mM MgSO4 and treated with 50 μg mL^-1^ RNase A (Roche) and 10 μg mL^-1^ DNase (Sigma-Aldrich) for additional 2 h at 37°C. Subsequently, 10 mM CaCl_2_ and 100 μg mL^-1^ trypsin (Sigma-Aldrich) were added and the mixture incubated at 37°C overnight. The next day 10 mM EDTA pH 7.4 were added. The solution was supplemented with SDS to a final concentration of 1 % (v/v) and boiled for 15 min in a bain-marie. This solution was cooled down to RT and centrifuged at 108,000 x g at RT for 45 min. The pellet was washed five times to remove any SDS traces. The isolated PG was dried in a SpeedVac (Thermo Scientific, Osterode, Germany) and stored at −20°C until further usage.

### Generation of *siiA* mutant alleles

Site-directed mutagenesis of low copy number vector p3187 harbouring siiA::HA under control of *P_sii_A* was performed by using the Q5 SDM kit according to the manufacturer instructions (NEB) (Table S9), with oligonucleotides (IDT, Munich, Germany) listed Table S7. The resulting plasmids were confirmed by DNA sequencing and introduced in *S*. Typhimurium strain MvP771 (Δ*siiA*::FRT, Table S8).

### *In-vitro* PG-binding assays

For *in vitro* binding assays, the proteins SiiA-PD, SiiA-R162A-PD, SiiA-R167L-PD, SiiA-S197E-PD and MotB-PD were dialyzed against buffer A (10 mM potassium phosphate pH 5.8) and/or buffer B (10 mM potassium phosphate 8.0). After buffer exchange, the protein variants were centrifuged at 16,100 x g for 15 min at 4°C in order to remove any precipitate. Isolated PG from *Salmonella* was resuspended and washed three times with buffer A or B. 0.2 mg of PG were mixed with 0.25 mg of purified protein in a 1.5 mL reaction tube. This was the input sample (sample IP). This mixture was incubated at RT for 1 h. The PG and PG-bound protein were pelleted via centrifugation at 16,100 x g for 10 min at 22°C and the supernatant collected in a separate tube. The pellet was re-suspended in 200 μL with the respective buffer. Unbound protein was removed from the insoluble PG by two additional centrifugation and resuspension steps, as described above. Samples for SDS-PAGE were taken from the resuspended pellet (samples P1, P2, P3…) and the supernatant (samples S1, S2, S3…) at each of the steps. The samples were analysed via SDS-PAGE and stained with Coomassie Blue. The band intensities were quantified with program ImageJ (https://imagej.nih.gov/ij/).

For analyses of PG binding in *S*. Typhimurium cells, assays were performed basically as described by Mizuno (1979) [60] (Table S8). The *S*. Typhimurium *siiA* deletion strain MvP771 harbouring plasmids for expression of WT *siiA*::HA or *siiA*-R162A::HA was grown overnight in LB, diluted 1:31 on LB and subcultured for 2.5 h for induction of SPI4 genes. Bacterial cells were harvested and resuspended in PBS, pH 7.4. Cells were disrupted by sonication using a Branson sonifier. The lysates were centrifuged for 60 min at 130,000 x g at 4°C and pellets were washed in PBS. Pellets were incubated in 900 μL extraction buffer (100 mM Tris-HCl, pH 7.3, 100 mM CaCl_2_, 10 % glycerol and 2 % SDS) at 34.6°C. After centrifugation for 130,000 x g for 1 h at 34.6°C, the supernatant was recovered and protein was precipitated by addition of TCA to a final concentration of 10 %. Precipitated protein was recovered by centrifugation at 16,100 x g for 46 min at 25°C. Pellets were washed with acetone, dried, and protein was dissolved in SDS-PAGE sample buffer. If required, pH was neutralized by addition of 1 M Tris-HCl, pH 10.0, and samples were denatured by boiling of 5 min.

### Infection of polarized epithelial cells

*S*. Typhimurium strains were grown over night in LB containing 50 μg/mL carbenicillin with aeration by continuous rotation in a roller drum at 37°C. Cultures were diluted 1:31 in LB and grown for 3.5 h at the same conditions in order to induce expression of SPI1 and SPI4 genes. Infection of the polarized epithelial cell line MDCK was performed as described before, and the amounts of internalized *Salmonella* were determined 1 h after infection [10].

### Measurement of intracellular pH

Plasmids were constructed using assembly cloning of PCR products [61]. All primers used for PCR are listed in Table S7. For the cloning of plasmids that allow co-expression of *siiAB, siiA*-D13N-B or *motAB* with pHluorin, pTAC-MAT-Tag-2 (Sigma-Aldrich) was PCR-amplified with primers pTAC-Gbs-for/rev (Table S9). As inserts, a tetracycline-inducible promoter was amplified from pWRG603 with primers TetR-Gbs-for/rev [10]. A codon-optimized version of R-pHluorin-M153R was synthesized (IDT) and amplified with primers R-pHluorin-tetR-Gbs-for/R-pHluorin-pTAC-Gbs-rev, and *siiAB, motAB* or *siiA*-D13N-*B* from *S*. Typhimurium genomic DNA or pWRG648 using primer pairs pTAC-SiiA-Gbs-for/SiiB462-tetR-Gbs-rev or pTAC-MotA-Gbs-for/MotB-pTAC-Gbs-rev [10, 34]. Plasmids containing further site-specific mutations in *siiA* (R162A) and *motB* (D33N) were generated by assembly cloning using primers listed in Table S7.

Overnight cultures of strains carrying pHluorin constructs were reinoculated at an OD_600_ of 0.15 in fresh LB broth containing 50 μg/mL carbenicillin (Carl Roth) and grown with aeration for 3.5 h at 37°C. Expression of *siiAB* or *motAB* together with R-pHluorin-M153R was induced after 2 h with 10 mM IPTG and 50 ng/mL anhydrotetracycline hydrochloride (Sigma-Aldrich). One mL of the suspension was centrifuged (8,000 x g, 5 min.), washed once with 1 mL 100 mM phosphate buffer pH 6.0 and finally resuspended in 500 μL of the same buffer. After 10-fold dilution fluorescence was determined with excitation at 410 and 470 nm and emission at 509 nm in a black 96-well plate (Nunc Thermo Fisher, Karlsruhe, Germany) using an Infinite M1000 plate reader (Tecan, Männedorf, Switzerland). The 410/470 nm excitation ratio was used to calculate the pH based on calibration curves. Calibration curves were generated with R-pHluorin-M153R expressing bacteria which were washed and resuspended in phosphate buffers of different pH ranging from 6.0 to 8.0. The protonophore carbonyl cyanide *m*-chlorophenyl hydrazone (CCCP, Sigma-Aldrich) was added to a final concentration of 30 μM and samples were incubated for 30 min. at 37°C. The ratio 410/470 nm was plotted against the pH-value and the resulting curve was fitted using a 4-parameter logistic function.

### Western blot

Bacterial cells were grown and *siiAB* expression was induced as described for intracellular pH measurements. An aliquot of the bacterial suspension was centrifuged (8,000 x g, 5 min.) and the pellet was resuspended in sample buffer (Roti-Load 1; Carl Roth) to normalize the volume to equal OD_600_. After heating (95°C, 5 min.) 10 μL were loaded onto 10 % polyacrylamide gels and separated in Tris-Tricine buffer according to manufacturer’s instructions (ProSieve; Lonza, Basel, Switzerland). The gels were wet blotted (Mini Trans-Blot Cell; Bio-Rad, Munich, Germany) onto polyvinylidene fluoride membranes (Immobilon-P, 0.45 μm; Merck-Millipore, Darmstadt, Germany). Membranes were probed using anti-HA or anti-SiiA and anti-SiiB primary antibodies (as indicated in the respective figures) [10] and horseradish peroxidase-coupled anti-rabbit secondary antibodies (Jackson ImmunoResearch, Ely, UK). Signals were visualized using a Chemi-Smart 3000 chemiluminescence system (Vilber Lourmat, Eberhardzell, Germany). Blot images were processed (marker overlay, tonal range, 16- to 8-bit conversion) using Photoshop CS6 (Adobe Systems, Munich, Germany).

### Accession numbers

Coordinates and structure factor amplitudes have been deposited with the protein data bank with accession number: 6QVP

## Acknowledgements

We would like to thank Manfred Weiss for help with data collection at beamline 14.1 at BESSY synchrotron and Max Kraner from the Biochemistry Division of the Department of Biology at Friedrich-Alexander University Erlangen-Nuremberg for mass spectrometry measurements.

## Abbreviations

Aa: Amino acid
CD: circular dichroism
IM: inner membrane
LR: linker region
MD: membrane domain
OM: outer membrane
PEM: periplasmic region essential for mobility
PG: peptidoglycan
PD: periplasmic domain
PMF: proton motive force
ppr: periplasmic region
rmsd: root mean square deviation
RT: room temperature
SPI: *Salmonella* pathogenicity island
T1SS: type one secretion system
T3SS-1: type three secretion system encoded by SPI1
T_M_: melting temperatures
WT: wild-type

## References

[1] B. Barlag, M. Hensel. The Giant Adhesin SiiE of Salmonella enterica. Molecules 20 (2015) 1134.

[2] G. Arya, R. Holtslander, J. Robertson, C. Yoshida, J. Harris, J. Parmley, A. Nichani, R. Johnson, C. Poppe. Epidemiology, Pathogenesis, Genoserotyping, Antimicrobial Resistance, and Prevention and Control of Non-Typhoidal Salmonella Serovars. Current Clinical Microbiology Reports 4 (2017) 43–53.

[3] J.A. Crump, M. Sjolund-Karlsson, M.A. Gordon, C.M. Parry. Epidemiology, Clinical Presentation, Laboratory Diagnosis, Antimicrobial Resistance, and Antimicrobial Management of Invasive Salmonella Infections. Clin Microbiol Rev 28 (2015) 901–937.

[4] R.G. Gerlach, N. Cláudio, M. Rohde, D. Jäckel, C. Wagner, M. Hensel. Cooperation of Salmonella pathogenicity islands 1 and 4 is required to breach epithelial barriers. Cellular Microbiology 10 (2008) 2364–2376.

[5] D.L. LaRock, A. Chaudhary, S.I. Miller. Salmonellae interactions with host processes. Nature Reviews Microbiology 13 (2015) 191–205.

[6] R.G. Gerlach, D. Jäckel, B. Stecher, C. Wagner, A. Lupas, W.-D. Hardt, M. Hensel. Salmonella Pathogenicity Island 4 encodes a giant non-fimbrial adhesin and the cognate type 1 secretion system. Cellular Microbiology 9 (2007) 1834–1850.

[7] E. Morgan, A.J. Bowen, S.C. Carnell, T.S. Wallis, M.P. Stevens. SiiE Is Secreted by the Salmonella enterica Serovar Typhimurium Pathogenicity Island 4-Encoded Secretion System and Contributes to Intestinal Colonization in Cattle. Infection and Immunity 75 (2007) 1524–1533.

[8] M.H. Griessl, B. Schmid, K. Kassler, C. Braunsmann, R. Ritter, B. Barlag, Y.-D. Stierhof, K.U. Sturm, C. Danzer, C. Wagner, T.E. Schaeffer, H. Sticht, M. Hensel, Y.A. Muller. Structural Insight into the Giant Ca2+-Binding Adhesin SiiE: Implications for the Adhesion of Salmonella enterica to Polarized Epithelial Cells. Structure 21 (2013) 741–752.

[9] B. Peters, J. Stein, S. Klingl, N. Sander, A. Sandmann, N. Taccardi, H. Sticht, R.G. Gerlach, Y.A. Muller, M. Hensel. Structural and functional dissection reveals distinct roles of Ca2+-binding sites in the giant adhesin SiiE of Salmonella enterica. PLOS Pathogens 13 (2017) e1006418.

[10] T. Wille, C. Wagner, W. Mittelstadt, K. Blank, E. Sommer, G. Malengo, D. Dohler, A. Lange, V. Sourjik, M. Hensel, R.G. Gerlach. SiiA and SiiB are novel type I secretion system subunits controlling SPI4-mediated adhesion of Salmonella enterica. Cell Microbiol 16 (2014) 161–178.

[11] J. Zhou, L.L. Sharp, H.L. Tang, S.A. Lloyd, S. Billings, T.F. Braun, D.F. Blair. Function of Protonatable Residues in the Flagellar Motor of Escherichia coli: a Critical Role for Asp 32 of MotB. Journal of Bacteriology 180 (1998) 2729–2735.

[12] S. Zhu, M. Takao, N. Li, M. Sakuma, Y. Nishino, M. Homma, S. Kojima, K. Imada. Conformational change in the periplamic region of the flagellar stator coupled with the assembly around the rotor. Proc Natl Acad Sci U S A 111 (2014) 13523–13528.

[13] E. Cascales, R. Lloubes, J.N. Sturgis. The TolQ-TolR proteins energize TolA and share homologies with the flagellar motor proteins MotA-MotB. Mol Microbiol 42 (2001) 795–807.

[14] V. Braun, S. Gaisser, C. Herrmann, K. Kampfenkel, H. Killmann, I. Traub. Energy-coupled transport across the outer membrane of Escherichia coli: ExbB binds ExbD and TonB in vitro, and leucine 132 in the periplasmic region and aspartate 25 in the transmembrane region are important for ExbD activity. Journal of Bacteriology 178 (1996) 2836–2845.

[15] A.A. Ollis, M. Manning, K.G. Held, K. Postle. Cytoplasmic membrane protonmotive force energizes periplasmic interactions between ExbD and TonB. Molecular Microbiology 73 (2009) 466–481.

[16] T. Minamino, K. Imada. The bacterial flagellar motor and its structural diversity. Trends Microbiol 23 (2015) 267–274.

[17] J.A. Wojdyla, E. Cutts, R. Kaminska, G. Papadakos, J.T.S. Hopper, P.J. Stansfeld, D. Staunton, C.V. Robinson, C. Kleanthous. Structure and Function of the Escherichia coli Tol-Pal Stator Protein TolR. Journal of Biological Chemistry 290 (2015) 26675–26687.

[18] E.R. Hosking, C. Vogt, E.P. Bakker, M.D. Manson. The Escherichia coli MotAB Proton Channel Unplugged. Journal of Molecular Biology 364 (2006) 921–937.

[19] S. Kojima, K. Imada, M. Sakuma, Y. Sudo, C. Kojima, T. Minamino, M. Homma, K. Namba. Stator assembly and activation mechanism of the flagellar motor by the periplasmic region of MotB. Molecular Microbiology 73 (2009) 710–718.

[20] R. De Mot, J. Vanderleyden. The C-terminal sequence conservation between OmpA-related outer membrane proteins and MotB suggests a common function in both Gram-positive and Gram-negative bacteria, possibly in the interaction of these domains with peptidoglycan. Molecular Microbiology 12 (1994) 333–334.

[21] T. Minamino, N. Terahara, S. Kojima, K. Namba. Autonomous control mechanism of stator assembly in the bacterial flagellar motor in response to changes in the environment. Mol Microbiol 109 (2018) 723–734.

[22] W. Vollmer. Chapter 6 - Peptidoglycan. In: Tang Y-W, Sussman M, Liu D, Poxton I, Schwartzman J, editors. Molecular Medical Microbiology (Second Edition). Boston: Academic Press; 2015. p. 105–124.

[23] J.S. Park, W.C. Lee, K.J. Yeo, K.-S. Ryu, M. Kumarasiri, D. Hesek, M. Lee, S. Mobashery, J.H. Song, S.I. Kim, J.C. Lee, C. Cheong, Y.H. Jeon, H.-Y. Kim. Mechanism of anchoring of OmpA protein to the cell wall peptidoglycan of the gram-negative bacterial outer membrane. The FASEB Journal 26 (2012) 219–228.

[24] L.M. Parsons, F. Lin, J. Orban. Peptidoglycan recognition by Pal, an outer membrane lipoprotein. Biochemistry 45 (2006) 2122–2128.

[25] A. Roujeinikova. Crystal structure of the cell wall anchor domain of MotB,, a stator component of the bacterial flagellar motor: Implications for peptidoglycan recognition. Proceedings of the National Academy of Sciences of the United States of America 105 (2008) 10348–10353.

[26] L. Slabinski, L. Jaroszewski, L. Rychlewski, I.A. Wilson, S.A. Lesley, A. Godzik. XtalPred: a web server for prediction of protein crystallizability. Bioinformatics 23 (2007) 3403–3405.

[27] C.S. Heilingloh, S. Klingl, C. Egerer-Sieber, B. Schmid, S. Weiler, P. Muhl-Zurbes, J. Hofmann, J.D. Stump, H. Sticht, M. Kummer, A. Steinkasserer, Y.A. Muller. Crystal Structure of the Extracellular Domain of the Human Dendritic Cell Surface Marker CD83. J Mol Biol 429 (2017) 1227–1243.

[28] S. Jones, J.M. Thornton. Principles of protein-protein interactions. Proc Natl Acad Sci U S A 93 (1996) 13–20.

[29] L. Holm, L.M. Laakso. Dali server update. Nucleic Acids Research 44 (2016) W351–W355.

[30] E.F. Pettersen, T.D. Goddard, C.C. Huang, G.S. Couch, D.M. Greenblatt, E.C. Meng, T.E. Ferrin. UCSF Chimera—A visualization system for exploratory research and analysis. Journal of Computational Chemistry25 (2004) 1605–1612.

[31] E.C. Meng, E.F. Pettersen, G.S. Couch, C.C. Huang, T.E. Ferrin. Tools for integrated sequence-structure analysis with UCSF Chimera. BMC Bioinformatics 7 (2006) 339.

[32] P.W. Rose, A. Prlic, C. Bi, W.F. Bluhm, C.H. Christie, S. Dutta, R.K. Green, D.S. Goodsell, J.D. Westbrook, J. Woo, J. Young, C. Zardecki, H.M. Berman, P.E. Bourne, S.K. Burley. The RCSB Protein Data Bank: views of structural biology for basic and applied research and education. Nucleic Acids Res 43 (2015) D345–356.

[33] S. Nakamura, N. Kami-ike, J.-i.P. Yokota, S. Kudo, T. Minamino, K. Namba. Effect of Intracellular pH on the Torque–Speed Relationship of Bacterial Proton-Driven Flagellar Motor. Journal of Molecular Biology 386 (2009) 332–338.

[34] Y.V. Morimoto, S. Kojima, K. Namba, T. Minamino. M153R Mutation in a pH-Sensitive Green Fluorescent Protein Stabilizes Its Fusion Proteins. PLOS ONE 6 (2011) e19598.

[35] H. Celia, N. Noinaj, S.D. Zakharov, E. Bordignon, I. Botos, M. Santamaria, T.J. Barnard, W.A. Cramer, R. Lloubes,S.K. Buchanan. Structural insight into the role of the Ton complex in energy transduction. Nature 538 (2016) 60–65.

[36] R. Dennehy, M. Romano, A. Ruggiero, Y.F. Mohamed, S.L. Dignam, C.M. Troncoso, M. Callaghan, M.A. Valvano, R. Berisio, S. McClean. The Burkholderia cenocepacia peptidoglycan-associated lipoprotein is involved in epithelial cell attachment and elicitation of inflammation. Cellular Microbiology 19 (2017).

[37] J. Kuriyan, D. Eisenberg. The origin of protein interactions and allostery in colocalization. Nature 450 (2007) 983–990.

[38] D.F. Evans, G. Pye, R. Bramley, A.G. Clark, T.J. Dyson, J.D. Hardcastle. Measurement of gastrointestinal pH profiles in normal ambulant human subjects. Gut 29 (1988) 1035–1041.

[39] C. Atuma, V. Strugala, A. Allen, L. Holm. The adherent gastrointestinal mucus gel layer: thickness and physical state in vivo. American Journal of Physiology-Gastrointestinal and Liver Physiology 280 (2001) G922–G929.

[40] S. Kojima, M. Takao, G. Almira, I. Kawahara, M. Sakuma, M. Homma, C. Kojima, K. Imada. The Helix Rearrangement in the Periplasmic Domain of the Flagellar Stator B Subunit Activates Peptidoglycan Binding and Ion Influx. Structure 26 (2018) 590–598.e595.

[41] C. The UniProt. UniProt: the universal protein knowledgebase. Nucleic Acids Res 45 (2017) D158–D169.

[42] W. Wang, B.A. Malcolm. Two-stage PCR protocol allowing introduction of multiple mutations, deletions and insertions using QuikChange Site-Directed Mutagenesis. BioTechniques 26 (1999) 680–682.

[43] F.W. Studier. Protein production by auto-induction in high-density shaking cultures. Protein Expression and Purification 41 (2005) 207–234.

[44] S.M. Kelly, T.J. Jess, N.C. Price. How to study proteins by circular dichroism. Biochimica et Biophysica Acta (BBA) - Proteins and Proteomics 1751 (2005) 119–139.

[45] B. Rupp. Biomolecular Crystallography: Principles, Practice, and Application to Structural Biology. 1st ed. New York: Garland Science; 2009.

[46] U. Mueller, R. Förster, M. Hellmig, F.U. Huschmann, A. Kastner, P. Malecki, S. Pühringer, M. Röwer, K. Sparta, M. Steffien, M. Ühlein, P. Wilk, M.S. Weiss. The macromolecular crystallography beamlines at BESSY II of the Helmholtz-Zentrum Berlin: Current status and perspectives. The European Physical Journal Plus 130 (2015) 141.

[47] W. Kabsch. XDS. Acta Crystallographica Section D 66 (2010) 125–132.

[48] K.M. Sparta, M. Krug, U. Heinemann, U. Mueller, M.S. Weiss. XDSAPP2.0. Journal of Applied Crystallography 49 (2016) 1085–1092.

[49] P. Evans. Scaling and assessment of data quality. Acta Crystallographica Section D 62 (2006) 72–82.

[50] T.C. Terwilliger, P.D. Adams, R.J. Read, A.J. McCoy, N.W. Moriarty, R.W. Grosse-Kunstleve, P.V. Afonine, P.H. Zwart, L.-W. Hung. Decision-making in structure solution using Bayesian estimates of map quality: the PHENIX AutoSol wizard. Acta Crystallographica Section D 65 (2009) 582–601.

[51] P.D. Adams, P.V. Afonine, G. Bunkoczi, V.B. Chen, I.W. Davis, N. Echols, J.J. Headd, L.-W. Hung, G.J. Kapral, R.W. Grosse-Kunstleve, A.J. McCoy, N.W. Moriarty, R. Oeffner, R.J. Read, D.C. Richardson, J.S. Richardson, T.C. Terwilliger, P.H. Zwart. PHENIX: a comprehensive Python-based system for macromolecular structure solution. Acta Crystallographica Section D 66 (2010) 213–221.

[52] P.V. Afonine, R.W. Grosse-Kunstleve, N. Echols, J.J. Headd, N.W. Moriarty, M. Mustyakimov, T.C. Terwilliger, A. Urzhumtsev, P.H. Zwart, P.D. Adams. Towards automated crystallographic structure refinement with phenix.refine. Acta Crystallographica Section D 68 (2012) 352–367.

[53] P. Emsley, B. Lohkamp, W.G. Scott, K. Cowtan. Features and development of Coot. Acta Crystallographica Section D 66 (2010) 486–501.

[54] T.C. Terwilliger, R.W. Grosse-Kunstleve, P.V. Afonine, N.W. Moriarty, P.H. Zwart, L.-W. Hung, R.J. Read, P.D. Adams. Iterative model building, structure refinement and density modification with the PHENIX AutoBuild wizard. Acta Crystallographica Section D 64 (2008) 61–69.

[55] E. Jurrus, D. Engel, K. Star, K. Monson, J. Brandi, L.E. Felberg, et al. Improvements to the APBS biomolecular solvation software suite. Protein Science 27 (2018) 112–128.

[56] D. Sitkoff, K.A. Sharp, B. Honig. Accurate Calculation of Hydration Free Energies Using Macroscopic Solvent Models. The Journal of Physical Chemistry 98 (1994) 1978–1988.

[57] W. Kabsch. A solution for the best rotation to relate two sets of vectors. Acta Crystallographica Section A 32 (1976) 922–923.

[58] E. Krissinel, K. Henrick. Inference of Macromolecular Assemblies from Crystalline State. Journal of Molecular Biology 372 (2007) 774–797.

[59] B. Glauner. Separation and quantification of muropeptides with high-performance liquid chromatography.Analytical Biochemistry 172 (1988) 451–464.

[60] T. Mizuno. A Novel Peptidoglycan-Associated Lipoprotein Found in the Cell Envelope of *Pseudomonas aeruginosa* and *Escherichia coli*. The Journal of Biochemistry 86 (1979) 991–1000.

[61] D.G. Gibson, L. Young, R.-Y. Chuang, J.C. Venter, C.A. Hutchison Iii, H.O. Smith. Enzymatic assembly of DNA molecules up to several hundred kilobases. Nature Methods 6 (2009) 343.

[62] G. Sheldrick. Experimental phasing with SHELXC/D/E: combining chain tracing with density modification.Acta Crystallographica Section D 66 (2010) 479–485.

[63] S. Li, T. Li, Y. Xu, Q. Zhang, W. Zhang, S. Che, R. Liu, Y. Wang, M. Bartlam. Structural insights into YfiR sequestering by YfiB in Pseudomonas aeruginosa PAO1. Sci Rep 5 (2015) 16915.

[64] M.L. Parker, S. Houston, C. Wetherell, C.E. Cameron, M.J. Boulanger. The Structure of Treponema pallidum Tp0624 Reveals a Modular Assembly of Divergently Functionalized and Previously Uncharacterized Domains. PLOS ONE 11 (2016) e0166274.

[65] R. Capelli, E. Matterazzo, M. Amabili, C. Peri, A. Gori, P. Gagni, M. Chiari, G. Lertmemongkolchai, M. Cretich, M. Bolognesi, G. Colombo, L.J. Gourlay. Designing Probes for Immunodiagnostics: Structural Insights into an Epitope Targeting Burkholderia Infections. ACS Infectious Diseases 3 (2017) 736–743.

[66] J. O’Neill, M. Xie, M. Hijnen, A. Roujeinikova. Role of the MotB linker in the assembly and activation of the bacterial flagellar motor. Acta Crystallographica Section D 67 (2011) 1009–1016.

[1] G.E. Crooks, G. Hon, J.M. Chandonia, S.E. Brenner. WebLogo: a sequence logo generator. Genome Res 14 (2004) 1188–1190.

