## Supplemental data for "Structural and functional characterisation of SiiA, an auxiliary protein from the SPI4-encoded type 1 secretion system from *Salmonella enterica*"

**Table S1. Melting temperatures analysed by CD-monitored thermal unfolding**

| **Protein** | **$T_M$ [°C]** | **pH** | **Figure** |
|---|---|---|---|
| **ppr-SiiA** | 76.8 +- 0.4 | 7.4 | S4g |
| **SiiA-PD** | 76.9 +-0.3 | 5.8 | S4a |
| **SiiA-PD** | 74.0 +-0.3 | 8.0 | S4a |
| **ppr-SiiA-R162A** | 86.1 +- 0.8 | 7.4 | S4h |
| **SiiA-R162A-PD** | 76.2 +- 0.7 | 5.8 | S4b |
| **SiiA-R162A-PD** | 75.2 +- 0.3 | 8.0 | S4b |
| **SiiA-R167L-PD** | 92.7 +- 2.8 | 5.8 | S4c |
| **SiiA-R167L-PD** | 90.4 +- 1.8 | 8.0 | S4c |
| **SiiA-S197E-PD** | 81.6 +- 0.6 | 5.8 | S4d |
| **SiiA-S197E-PD** | 75.9 +- 0.4 | 8.0 | S4d |
| **MotB-PD** | 63.2 +- 0.1 | 5.8 | S4e |
| **MotB-PD** | 63.2 +- 0.2 | 8.0 | S4e |

**Table S2. Analytical size exclusion chromatography results**

| **Protein** | **MW [kDa]** | **$V_e$ [mL]** | |
|---|---|---|---|
| **Dextranblue** | | 7.856 (= $V_0$) | a [a] |
| **Coalbumin** | 75 | 9.2 | b [a] |
| **Carboanhydrase** | 29 | 11.4 | c [a] |
| **RNase A** | 13.7 | 12.9 | d [a] |
| **Aprotein** | 6.5 | 14.9 | e [a] |
| **ppr-SiiA** | 44.84 [b] | 10.34 | |
| **SiiA-PD** | 17.31 [b] | 12.52 | |
| **ppr-SiiA-R162A** | 44.34 [b] | 10.37 | |
| **SiiA-R162-PD** | 17.07 [b] | 12.56 | |

[a] The elution volumes of the protein standards are marked with the letters a, b, c, d and e in Fig. S1b,c).

[b] Apparent molecular weights (MW) calculated *via* a least squared fit of a trendline based upon the elution volume ($V_e$). Formular: y = -0.189 * x + 3.609 ($R^2$ = 0.999).

**Table S3. Peak assignment in the anomalous difference density map**

| Number | Peak height (peak intensity/standard deviation) | Corresponding aa[a] |
|---|---|---|
| **1** | 18.7 | MSE, E.172 |
| **2** | 18.1 | MSE, F.186 |
| **3** | 17.9 | MSE, D.186 |
| **4** | 17.0 | MSE, E.186 |
| **5** | 17.0 | MSE, E.172 |
| **6** | 16.5 | MSE, F.172 |
| **7** | 16.0 | MSE, C.148/A |
| **8** | 15.5 | MSE, B.148/A |
| **9** | 15.4 | MSE, D.148/A |
| **10** | 15.0 | MSE, A.148 |
| **11** | 13.8 | MSE, F.148 |
| **12** | 13.6 | MSE, E.148 |
| **13** | 13.3 | MSE, B.186 |
| **14** | 13.0 | MSE, C.186 |
| **15** | 12.8 | MSE, A.186 |
| **16** | 11.3 | MSE, A.172 |
| **17** | 11.2 | MSE, C.172 |
| **18** | 9.5 | MSE, B.172 |
| **19** | 6.5 | MSE, B.148/B |
| **20** | 5.8 | MSE, C.148/B |
| **21** | 5.4 | MSE, D.148/B |

[a] Residues are numbered as follows: residue, chain.residue number/alternative conformation. MSE = Seleno-methionine residue.

**Table S4. $C_\alpha$-rmsd values between SiiA-PD monomers**

| | SiiA_A[a] | SiiA_B | SiiA_C | SiiA_D | SiiA_E | SiiA_F |
|---|---|---|---|---|---|---|
| **SiiA_A** | - | 1.0[b] | 0.1 | 1.0 | 0.2 | 0.9 |
| **SiiA_B** | | - | 1.0 | 0.2 | 1.0 | 0.2 |
| **SiiA_C** | | | - | 1.0 | 0.1 | 0.9 |
| **SiiA_D** | | | | - | 1.0 | 0.3 |
| **SiiA_E** | | | | | - | 0.9 |
| **SiiA_F** | | | | | | - |

[a] White- and grey-shaded protein chain entries belong to group 1 and 2, respectively.
[b] Values calculated with program LSQKAB [1].

**Table S5. $C_\alpha$-rmsd values obtained upon comparing SiiA-PD dimers**

| | SiiA_AB | SiiA_BA | SiiA_CD | SiiA_DC | SiiA_EF | SiiA_FE |
|---|---|---|---|---|---|---|
| **SiiA_AB** | - | 2.6[a, b] | 0.2 | 2.6 | 0.2 | 2.6 |
| **SiiA_BA** | | - | 2.6 | 0.2 | 2.6 | 0.2 |
| **SiiA_CD** | | | - | 2.6 | 0.3 | 2.6 |
| **SiiA_DC** | | | | - | 2.6 | 0.3 |
| **SiiA_EF** | | | | | - | 2.7 |
| **SiiA_FE** | | | | | | - |

[a] Values calculated with program LSQKAB [1].
[b] The SiiA-AB dimer is compared to the SiiA-BA dimer such that chain A is superimposed onto chain B and at the same time chain B onto chain A. This applies accordingly to all dimer comparisons.

**Table S6. Peptidoglycan co-isolation quantification**

| | Signal intensity anti HA | Signal intensity anti OmpA[a] | Ratio SiiA/OmpA |
|---|---|---|---|
| WT SiiA | 30,774 | 14,594 | 2.11 |
| SiiA-R162A | 13,459 | 14,203 | 0.95 |
| Ratio WT /mut | 2.29 | 1.02 | |

[a] Antibody staining against WT OmpA was used as a control to normalize the precipitated PG amount.

**Table S7. Primers used for generation of expression constructs, site-directed mutagenesis, and pH sensors**

| Primer | Sequence (5’ -> 3’) | Usage |
|---|---|---|
| **R162A-fwd** | GACATCTCTTTCTCTGACTCTCTAGCACTGGGATATGAAG | Recombinant protein expression |
| **R162A-rev** | GTTCATATCCCAGTGCTAGAGAGTCAGAGAAAGAGATGTC | Recombinant protein expression |
| **R167L-fwd** | GGGATATGAActgGGAATTATTTTG | Recombinant protein expression |
| **R167L-rev** | AGTCGTAGAGAGTCAGAG | Recombinant protein expression |
| **S197E-fwd** | AAGTACAACGgaaAAAGCTATTATCACGAC | Recombinant protein expression |
| **S197E-rev** | GATGCTGCGGAGTTAACAC | Recombinant protein expression |
| **R120A-For** | GTATCATGGCgcgCTGAGAAGCTTTTC | SDM[a] |
| **R120A-Rev** | GTAATAACAAGCTCATTTTTTGTAG | SDM |
| **D159A-For** | CTCTTTCTCTgccTCTCTACGAC | SDM |
| **D159A-Rev** | ATGTCTGCCTGAGGAATAAC | SDM |
| **R162A-For** | TGACTCTCTAgcaCTGGGATATGAACG | SDM |
| **R162A-Rev** | GAGAAAGAGATGTCTGCC | SDM |
| **L163A-For** | CTCTCTACGAgcgGGATATGAACGG | SDM |
| **L163A-Rev** | TCAGAGAAAGAGATGTCTG | SDM |
| **R167L-For** | GGGATATGAActgGGAATTATTTTG | SDM |
| **R167L-Rev** | AGTCGTAGAGAGTCAGAG | SDM |
| **S197E-For** | AAGTACAACGgaaAAAGCTATTATCACGAC | SDM |
| **S197E-Rev** | GATGCTGCGGAGTTAACAC | SDM |
| **MotB-D32N-Gbs-for** | GGAAAATTGCCTACGCCAATTTTATGACGGCGATGATGGC | Intracellular pH measurements |
| **MotB-D32N-Gbs-rev** | GCCATCATCGCCGTCATAAAATTGGCGTAGGCAATTTTCC | Intracellular pH measurements |
| **MotB-pTAC-Gbs-rev** | AACCAAGATGTCGAGTTAACCACCCATCGATCACCTCGGTTCCGCTTTTG | Intracellular pH measurements |
| **pTAC-Gbs-for** | CGATCTCGACGAGTGAGAGAAG | Intracellular pH measurements |
| **pTAC-Gbs-rev** | CATATTATATCTCCTGTGTGAAATTGTTATCC | Intracellular pH measurements |
| **pTAC-MotA-Gbs-** | GATAACAATTTCACACAGGAGATATAATATG | Intracellular pH |

| | | |
|---|---|---|
| **for** | CTTATCTTATTAGGTTACCTG | measurements |
| **pTAC-SiiA-Gbs-for** | GATAACAATTTCACACAGGAGATATAATATG GAAGACGAAAGTAATCCG | Intracellular pH measurements |
| **pWRG692-Gbs-for** | CCATGGATCGATAGCTGGTC | Intracellular pH measurements |
| **R-pHluorin-pTAC-Gbs-rev** | TGAAAATCTTCTCTCACTCGTCGAGATCGTTA TTATTTATACAGTTCATC | Intracellular pH measurements |
| **R-pHluorin-pTAC-Gbs-rev** | TGAAAATCTTCTCTCACTCGTCGAGATCGTTA TTATTTATACAGTTCATC | Intracellular pH measurements |
| **R-pHluorin-tetR-Gbs-for** | ATAGAGAAAGATGGCAAGAGGAGGATATCA TGTCTAAAGGCGAAGAACTG | Intracellular pH measurements |
| **R-pHluorin-tetR-Gbs-for** | ATAGAGAAAGATGGCAAGAGGAGGATATCA TGTCTAAAGGCGAAGAACTG | Intracellular pH measurements |
| **SiiA-R162A-pTAC-Gbs-for** | ATCTCTTTCTCTGACTCTCTAGCGCTGGGATA TGAACGGGGAATTATTTTGATGAAAGAG | Intracellular pH measurements |
| **SiiA-R162A-pTAC-Gbs-rev** | CTCTTTCATCAAAATAATTCCCCGTTCATATC CCAGCGCTAGAGAGTCAGAGAAAGAGAT | Intracellular pH measurements |
| **SiiB462-tetR-Gbs-rev** | GTAACCAAGATGTCGAGTTAACCACCCATCG ATTAATCTTCATTTTTTTCCTCCTTG | Intracellular pH measurements |
| **TetR-Gbs-for** | TCGATGGGTGGTTAACTCGAC | Intracellular pH measurements |
| **TetR-Gbs-rev** | GATATCCTCCTCTTGCCATC | Intracellular pH measurements |

[a] SDM, site-directed mutagenesis.

**Table S8. Bacterial strains used in this study**

| Designation | relevant genotype | Source/reference |
|---|---|---|
| *Salmonella enterica* serovar Typhimurium | | |
| **NCTC 12023** | WT | NCTC Colindale, lab collection |
| **MvP771** | Δ*siiA*::FRT | [2] |
| **MvP103** | Δ*sseC*::*aphT*, $Kan^R$ | [3] |
| *Escherichia coli* | | |
| **OneShot Mach1-$T1^R$** | F-φ80(*lacZ*)ΔM15 Δ*lacX*74 *hsdR*(rK-mK+) Δ*recA*1398 *endA*1 *tonA* | Sigma-Aldrich |
| **XL10Gold** | F´ *proA*$^+$*B*$^+$ *lacI*$^q$ Δ*(lacZ)M15 zzf::Tn10* ($Tet^R$)*/fhuA2*Δ*(argF-lacZ)U169 phoA glnV44 Φ80Δ(lacZ)M15 gyrA96 recA1 relA1 endA1 thi-1 hsdR17* | NEB |
| **BL21(DE3)** | $F^–$ *ompT hsdS*$_B$ ($r_B^–$ $m_B^–$) *gal dcm* (DE3) | Novagen |

**Table S9. Plasmids strains used in this study**

| Designation | relevant genotype | Source/reference |
|---|---|---|
| **pWSK29** | low copy number vector, $Amp^R$ | [4] |
| **p3187** | $P_{siiA}$ *siiA*::HA in pWSK29 | [5] |
| **p4478** | $P_{siiA}$ *siiA*::HA R120A | This study |
| **p4479** | $P_{siiA}$ *siiA*::HA D159A | This study |
| **p4480** | $P_{siiA}$ *siiA*::HA R162A | This study |
| **p4481** | $P_{siiA}$ *siiA*::HA L163A | This study |
| **p4482** | $P_{siiA}$ *siiA*::HA R167A | This study |
| **p4483** | $P_{siiA}$ *siiA*::HA S197E | This study |
| **pTAC-MAT-Tag-2** | *tac* promoter, $Amp^R$ | Sigma-Aldrich |
| **pWRG603** | $P_{tetA}$::*siiF*-G500E::*scfp3a*<br>$P_{tetA}$::*siiB*::*syfp2*, $Amp^R$ | [2] |
| **pWRG850** | $P_{tac}$-*siiAB*-*tetR*-$P_{tetA}$-*pHluorin*-M153R, $Amp^R$ | This study |
| **pWRG851** | $P_{tac}$-*siiA*-D13N-*B*-*tetR*-$P_{tetA}$-*pHluorin*-M153R, $Amp^R$ | This study |
| **pWRG855** | $P_{tac}$-*motAB*-*tetR*-$P_{tetA}$-*pHluorin*-M153R, $Amp^R$ | This study |
| **pWRG862** | $P_{tac}$-*motAB*-D32N-*tetR*-$P_{tetA}$-*pHluorin*-M153R, $Amp^R$ | This study |
| **pWRG898** | $P_{tac}$-*siiA*-R162A-*B*-*tetR*-$P_{tetA}$-*pHluorin*-M153R, $Amp^R$ | This study |
| **pET-15B** | Expression vector, $Amp^R$ | Novagen |
| **pBT127** | pET-15B, *ppr-siiA* | This study |
| **pBT380** | pET-15B, *ppr-siiA*-R162A | This study |
| **pBT427** | pET-15B, *motB*-PD | This study |
| **pBT444** | pET-15B, *ppr-siiA*-R167L | This study |
| **pBT445** | pET-15B, *ppr-siiA*-S197E | This study |

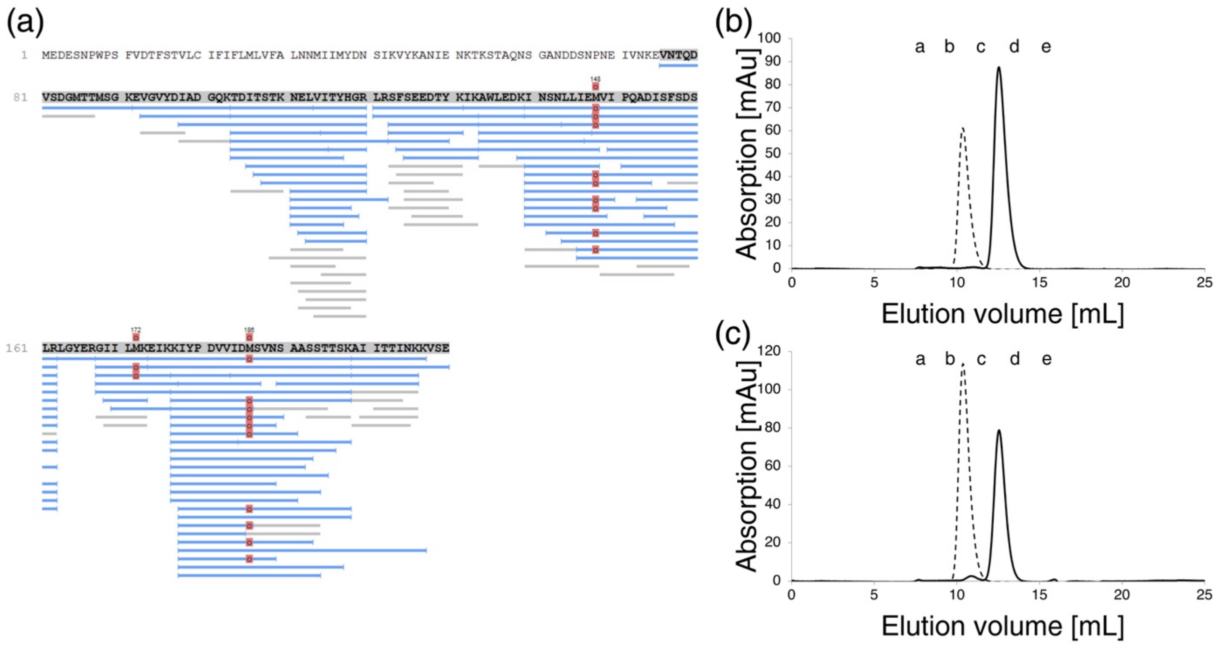


**Fig. S1. Mass spectrometry sequence mapping and analytical gel-filtration analysis of SiiA-PD.**

(a) Analysis of tryptic fragments of SiiA-PD (blue lines) by means of ESI mass spectrometry. Grey lines indicate observed fragments below the quality score (false discovery rate <20 %). Red dots mark oxidized methionines. Good sequence coverage is observed from residue Thr104 to Lys206. However, additional fragments are observed beyond this range. The analysis does not allow for the unambiguous determination of the exact starting and ending residue of the protease-generated SiiA-PD fragment. (b) Analytical size exclusion chromatography analysis of ppr-SiiA (broken line) and SiiA-PD (black line). (c) Analytical size exclusion chromatography analysis of ppr-SiiA-R162A (broken line) and SiiA-R162A-PD (black line). In panels (b) and (c) the labels a, b, c, d and e indicate the elution volumes of the protein standards listed in Table S2.

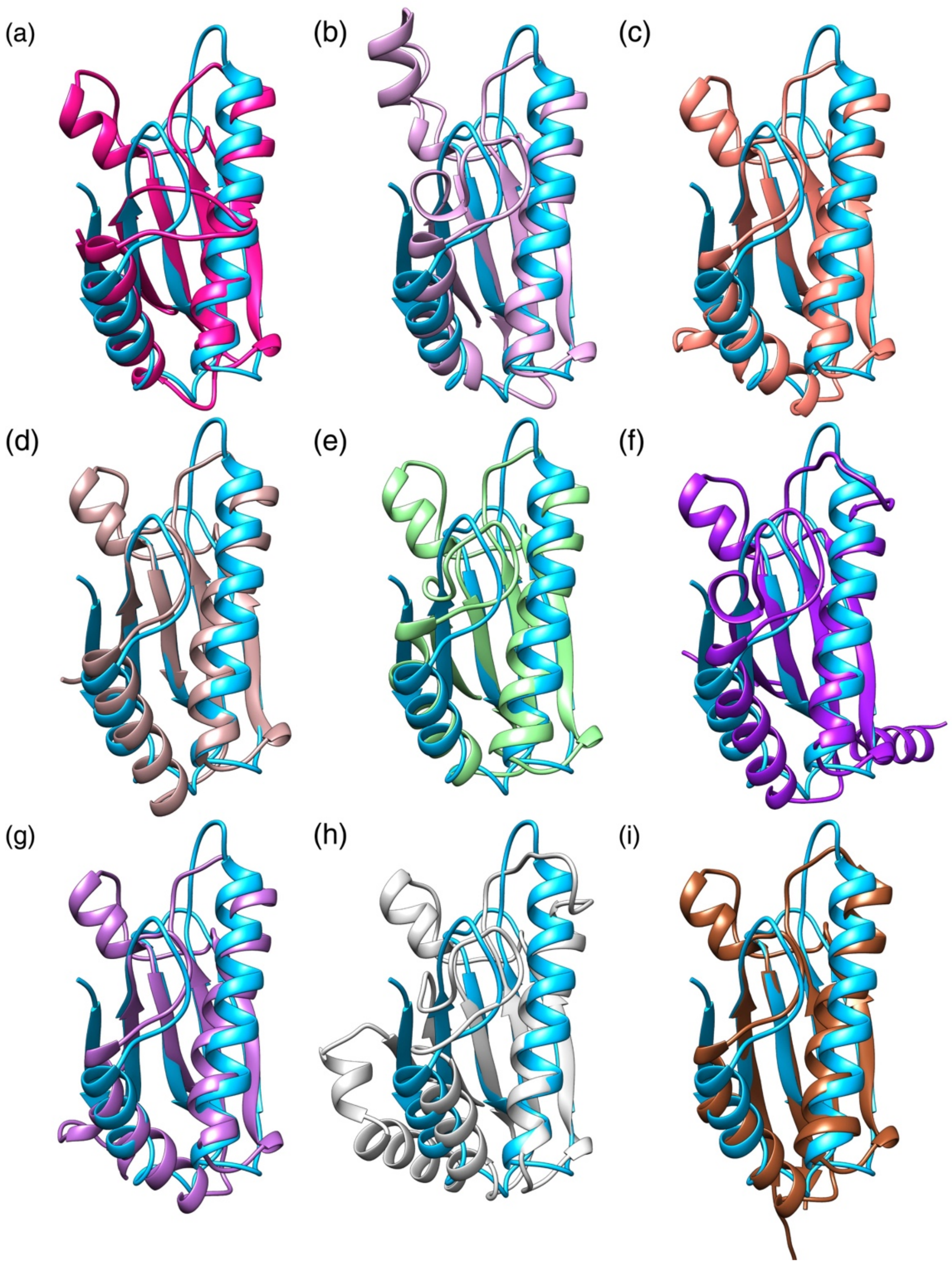


**Fig. S2. Structure comparison of SiiA-PD with various OmpA-like PG-binding domains.** Structural homologues of SiiA-PD identified with the DALI server [6] (see also Table 2 in main text): SiiA-PD (light blue) in comparison to (a) PAL from *T. pallidum* (PDB-ID: 5IJR, [7]), (b) OmpA from *K. pneumoniae* (PDB-ID: 5NHX), (c) PAL from *B. cenocepacia* (PDB-ID: 5N2C), (d) YifB from *P.aeruginosa* (PDB-ID: 4ZHW), (e) TagL from *E.coli* (PDB-ID: 5M38), (f) MotB from *H. pylori* (PDB-ID: 3S02), (g) PAL from *B. cenocepacia* (PDB-ID: 5LKW), (h) PomB from *V. alginolyticus* (PDB-ID: 3WPW) and (i) OmpA from *A. baumannii* (PDB-ID: 4G88).

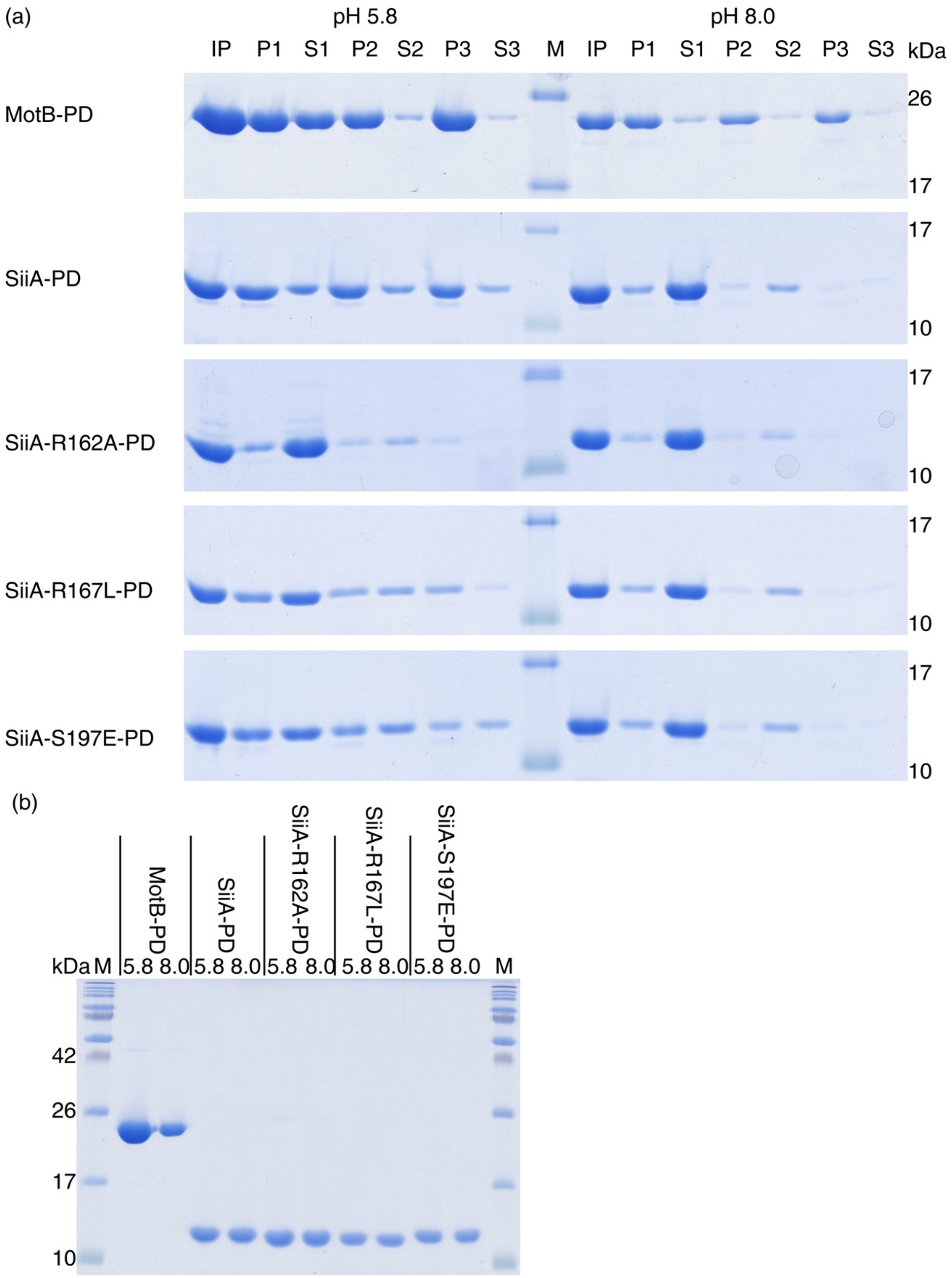
(a)
pH 5.8
pH 8.0
IP P1 S1 P2 S2 P3 S3 M IP P1 S1 P2 S2 P3 S3 kDa
MotB-PD
SiiA-PD
SiiA-R162A-PD
SiiA-R167L-PD
SiiA-S197E-PD
26
17
10
(b)
MotB-PD
SiiA-PD
SiiA-R162A-PD
SiiA-R167L-PD
SiiA-S197E-PD
kDa M 5.8 8.0 5.8 8.0 5.8 8.0 5.8 8.0 5.8 8.0 M
42
26
17
10

**Fig. S3. PG binding of MotB-PD, SiiA-PD, SiiA-R162A-PD, SiiA-R167L-PD and SiiA-S197E-PD.**

(a) PG pulldown assays performed at pH 5.8 (left side) and pH 8.0 (right side) with the different protein variants and analysed by SDS-PAGE. Input (IP), pellet (P1-3) and supernatant (S1-3) fractions, as well as the relevant marker bands are indicated. Shown are the original electrophoretic gels from which the bands that are displayed individually in Fig. 5 of the main text have been extracted. (b) SDS-PAGE analysis of the input samples of the protein variants used in (a). The molecular weights of selected marker bands are reported on the left side of the gel.

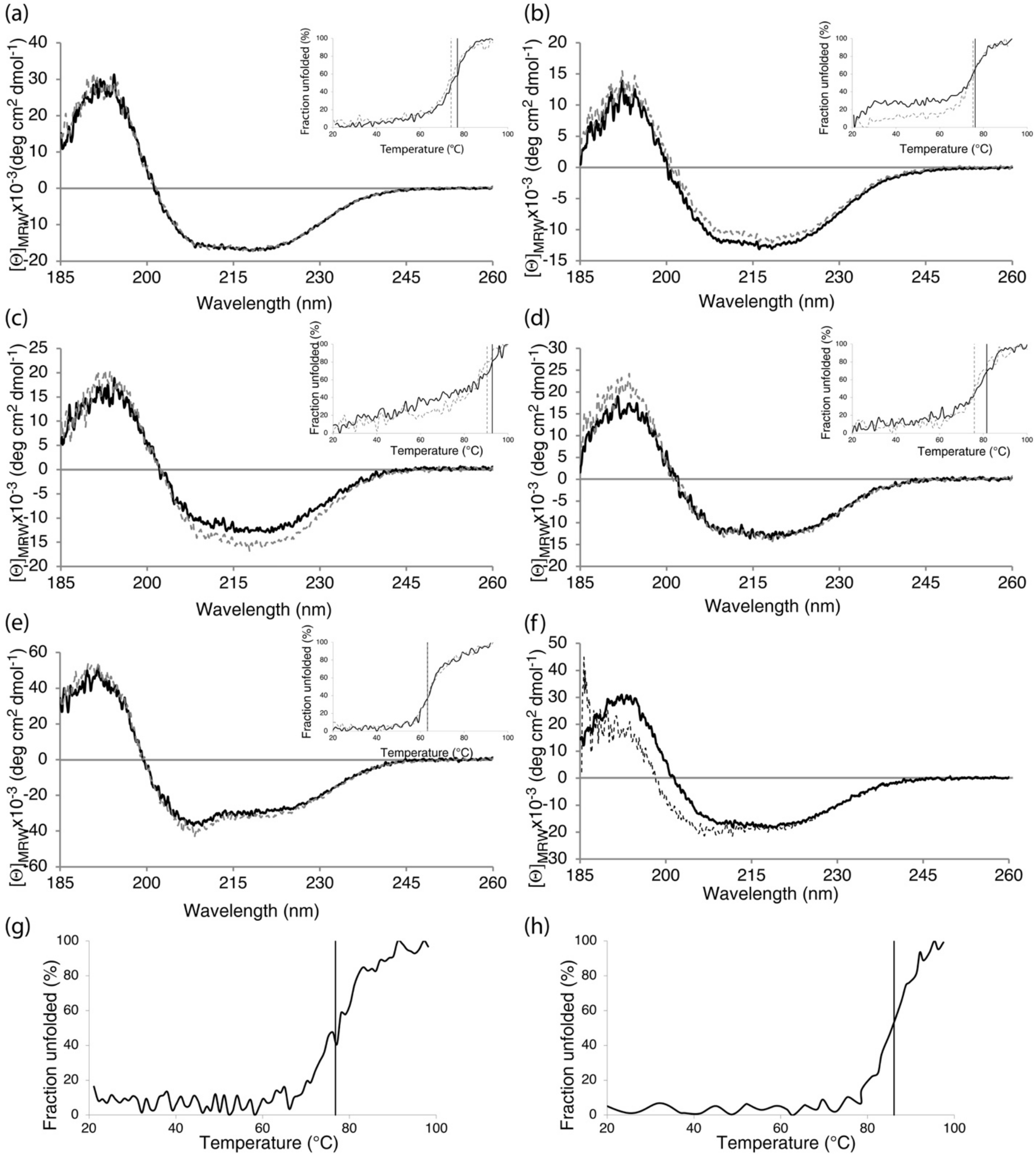


**Fig. S4. Structural integrity of SiiA variants and of MotB-PD probed by CD spectroscopy.** CD spectra of (a) SiiA-PD, (b) SiiA-R162A-PD, (c) SiiA-R167L-PD, (d) SiiA-S197E-PD and (e) MotB-PD fragments. Insets display the CD-monitored thermal denaturation of the respective proteins. In panels (e) to (f) contiguous black lines and dashed grey lines show the proteins in 10 mM potassium phosphate-based buffers at pH 5.8 and 8.0, respectively. (f) CD spectra of SiiA-

R162A-PD (black line) and ppr-SiiA-R162A (grey dashed line) recorded at pH 7.4. (g) and (h) show the thermal denaturation at pH 7.4 of ppr-SiiA and ppr-SiiA-R162A, respectively. The observed melting temperatures are summarized in Table S1 and marked with vertical lines in all panels.

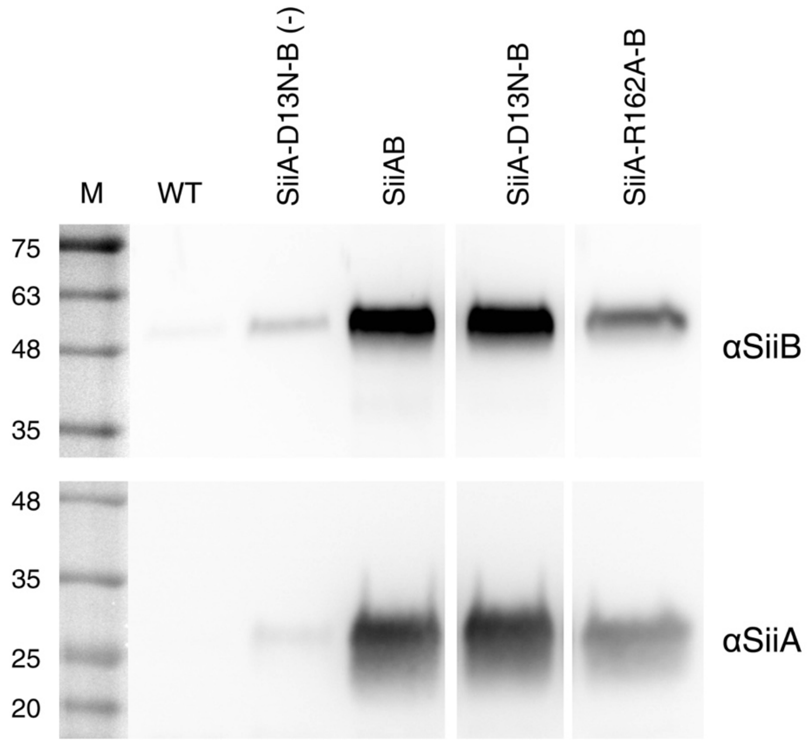


**Fig. S5. Inducible expression of *siiAB*.**

*S*. Typhimurium NCTC 12023 WT bacteria were transformed with plasmids allowing for co-production of SiiAB with R-pHluorin-M153R. Bacteria were subcultured with or without (-) the addition of 10 mM IPTG to induce expression of the indicated SiiAB complexes as described for intracellular pH measurements. The samples were separated in a 10% SDS polyacrylamide gel and blotted. The proteins were detected by specific antibodies against SiiA and SiiB. WT bacteria without plasmid served as control. One Western blot from three independent experiments with similar results is shown. M = protein marker with molecular sizes in kDa as indicated.

### Additional references


[1] W. Kabsch. A solution for the best rotation to relate two sets of vectors. Acta Crystallographica Section A 32 (1976) 922-923.

[2] T. Wille, C. Wagner, W. Mittelstadt, K. Blank, E. Sommer, G. Malengo, D. Dohler, A. Lange, V. Sourjik, M. Hensel, R.G. Gerlach. SiiA and SiiB are novel type I secretion system subunits controlling SPI4-mediated adhesion of Salmonella enterica. Cell Microbiol 16 (2014) 161-178.

[3] E. Medina, P. Paglia, T. Nikolaus, A. Müller, M. Hensel, C.A. Guzmán. Pathogenicity Island 2 Mutants of <em>Salmonella typhimurium</em> Are Efficient Carriers for Heterologous Antigens and Enable Modulation of Immune Responses. Infection and Immunity 67 (1999) 1093-1099.

[4] W. Rong Fu, S.R. Kushner. Construction of versatile low-copy-number vectors for cloning, sequencing and gene expression in Escherichia coli. Gene 100 (1991) 195-199.

[5] R.G. Gerlach, D. Jäckel, B. Stecher, C. Wagner, A. Lupas, W.-D. Hardt, M. Hensel. Salmonella Pathogenicity Island 4 encodes a giant non-fimbrial adhesin and the cognate type 1 secretion system. Cellular Microbiology 9 (2007) 1834-1850.

[6] L. Holm, L.M. Laakso. Dali server update. Nucleic Acids Research 44 (2016) W351-W355.

[7] P.W. Rose, A. Prlic, C. Bi, W.F. Bluhm, C.H. Christie, S. Dutta, R.K. Green, D.S. Goodsell, J.D. Westbrook, J. Woo, J. Young, C. Zardecki, H.M. Berman, P.E. Bourne, S.K. Burley. The RCSB Protein Data Bank: views of structural biology for basic and applied research and education. Nucleic Acids Res 43 (2015) D345-356.
